# Identification of RNA base pairs and complete assignment of nucleobase resonances by proton-detected solid-state NMR spectroscopy at 100 kHz MAS

**DOI:** 10.1101/2021.07.11.450890

**Authors:** Philipp Innig Aguion, John Kirkpatrick, Teresa Carlomagno, Alexander Marchanka

## Abstract

Knowledge of RNA structure, either in isolation or in complex, is fundamental to understand the mechanism of cellular processes. Solid-state NMR (ssNMR) is applicable to high molecular-weight complexes and does not require crystallization; thus, it is well-suited to study RNA as part of large multicomponent assemblies. Recently, we solved the first structures of both RNA and an RNA–protein complex by ssNMR using conventional ^13^C- and ^15^N-detection. This approach is limited by the severe overlap of the RNA peaks together with the low sensitivity of multidimensional experiments. Here, we overcome the limitations in sensitivity and resolution by using ^1^H-detection at fast MAS rates. We develop experiments that allow the identification of complete nucleobase spin-systems together with their site-specific base pair pattern using sub-milligram quantities of one uniformly labelled RNA sample. These experiments provide rapid access to RNA secondary structure by ssNMR in protein–RNA complexes of any size.

## Introduction

Ribonucleic acids (RNA) perform a multitude of cellular functions, including the regulation of gene expression. The structures of the RNAs involved in these functions, and of their complexes with proteins, shed light on the molecular mechanisms of the corresponding cellular processes and facilitate intervention in a disease context. Nuclear magnetic resonance (NMR) spectroscopy in solution plays an important role in studying RNA structure, as it can handle well the inherent flexibility exhibited by many RNAs.^[1,2]^ However, due to the strong dependency of the resonance linewidths on the rate of rotational diffusion, solution-state NMR can be applied straightforwardly only to molecules of moderate size, typically up to molecular-weights (MW) of ~100 and 40 kDa for proteins and nucleic acids, respectively. Even for RNAs larger than 20 kDa, tailored labeling schemes, such as segmental or nucleotide specific ^13^C/^15^N/^2^H-labelling, are required to alleviate spectral overlap and reduce line-broadening.^[3–5]^ Recently, the Summers laboratory^[6]^ has used ^2^H-edited NMR to probe the three-dimensional structure of the 42 kDa HIV-1 ^Cap^1G-L^TPUA^ RNA; the study required a total of 16 nucleotide-type specifically labelled samples, demonstrating that the application of solution-state NMR to RNAs of MW ≥ 40 kDa remains challenging.

In solid-state NMR (ssNMR) the linewidth does not increase with the molecular size. As a consequence, the technique can be applied to particles of any size, provided that the sensitivity is sufficiently high and that peak-crowding does not become a limiting factor. For medium-sized RNAs embedded in large complexes, peak-crowding is rarely a problem, as typically only the RNA of interest is visible in isotope-edited NMR spectra.^[7,8]^ However, high-MW particles lead to low sensitivity, simply because the number of copies that can be packed in a given volume is inversely proportional to the particle size. Sensitivity has been improved by optimizing both the hardware and the experimental design, particularly with the recent development of ultra-fast MAS (magic-angle spinning) probe-heads that have allowed the advent of ^1^H-detection in biomolecular solid-state NMR.^[9–14]^

Over the last decade, we have developed experiments to assign ^13^C and ^15^N resonances of RNA bound to proteins. We demonstrated the methodology using the 8.5-kDa 26mer box C/D RNA in complex with the ~13.5 kDa protein L7Ae from *Pyrococcus furiosus* (*Pf*) and determined its structure by ssNMR (Fig. 1 a, b).^[7,8,15–17]^ In this work, we used ^15^N- and ^13^C-detection, which limited the sensitivity of the experiments and prevented acquisition of three-dimensional spectra. In addition, at the achievable MAS rate of 20 kHz, the ^15^N- and ^13^C-linewidths were such as to lead to significant peak-overlap, especially in the regularly structured A-form helical regions. As a consequence, solving the structure of the RNA required eight different samples with single and double nucleotide-type selective labelling.

**Figure 1.**
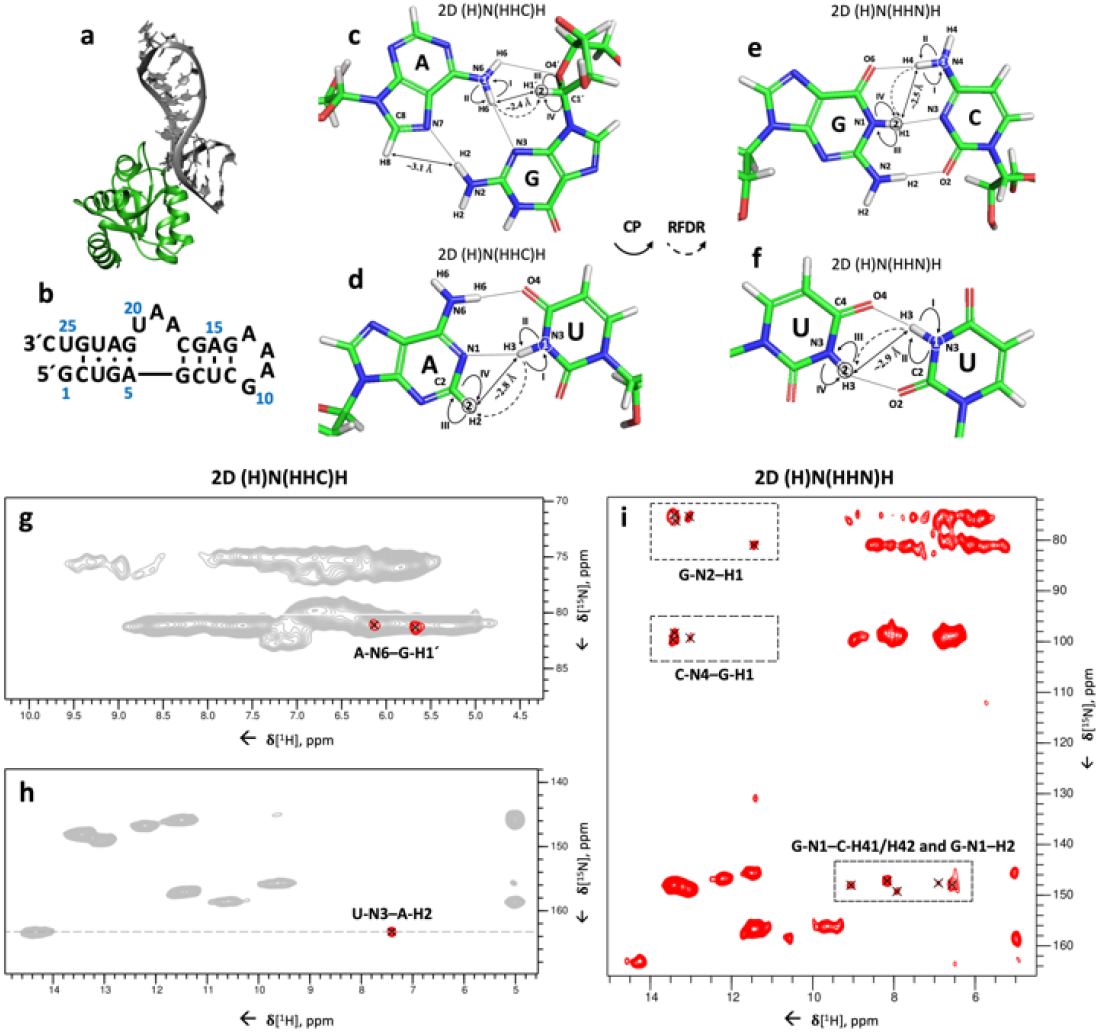
Identification of base pairs in the 26mer box C/D RNA. (**a**) Structure of the *Pf* 26mer box C/D RNA (grey) bound to the *Pf* L7Ae protein (lime green), determined by ssNMR using paramagnetic-relaxation-enhancement and chemical-shift-perturbation data (PDB entry 6TPH).^[15]^ (**b**) Sequence and secondary-structure of the *Pf* 26mer box C/D RNA. (**c-f**) Magnetization transfer schemes of the 2D (H)N(HHC)H (c-d) and 2D (H)N(HHC)H (e-f) experiments shown for G:A (**c**), A:U (**d**), G:C (**e**) and U:U (**f**) base pairs. Encircled numbers indicate the chemical-shift evolution times (*t*_*1*_ & *t*_*2*_) corresponding to the two spectral dimensions, and the roman numerals indicate the CP transfer periods. Inter-strand proton-proton distances involved in magnetization transfer via ^1^H-^1^H RFDR across the base pair are labelled. (**g-h**) Amino (**g**) and imino (**h**) regions from the 2D (H)N(HHC)H spectrum of the 26mer box C/D RNA in complex with L7Ae showing inter-strand A-N6–G-H1’ **(g)** and U-N3–A-H2 **(h)** cross-peaks specific for G:A and A:U base pairs, respectively. For reference, the red contours of the 2D (H)N(HHC)H spectrum are overlaid onto the 2D ^1^H-^15^N CP-HSQC spectrum (in grey). (**i**) 2D (H)N(HHC)H spectrum showing inter-strand C-N4–G-H1 and G-N1–C-H41/H42 cross-peaks as well as intra-residue G-N1–H2 and G-N2–H1 cross-peaks. The experimental and processing parameters are given in Table S1–3; pulse sequences and phase cycles are given in Fig. S1a and b. The ^1^H-^1^H RFDR mixing time was either 0.48 ms (2D (H)N(HHN)H) or 0.4 ms (2D (H)N(HHC)H). All spectra were recorded on a Bruker Avance III HD spectrometer at a ^1^H field-strength of 850 MHz, a MAS rate of 100 kHz, and a temperature of 275 K, using a 0.81-mm MAS probe-head developed by the Samoson group (https://www.nmri.eu/).

^1^H-detected ssNMR in combination with high-speed MAS combines the high sensitivity of ^1^H detection with sharp and MW-independent linewidths. To date, ^1^H-detection has been applied mostly to proteins,^[9–13]^ with ^1^H-detected ssNMR of RNA remaining somewhat of a challenge, due to the low proton density in the nucleobases and the poor ^1^H chemical-shift dispersion in the ribose ring. Nonetheless, the improved linewidth and sensitivity of ^1^H-detected ssNMR at a MAS rate of 100 kHz have allowed us to measure a range of three-dimensional spectra of uniformly ^13^C,^15^N-labelled 26mer box C/D RNA in complex with L7Ae,^[14]^ such as HCN-like^[18–20]^ experiments.

NMR spectroscopy, both in solution- and in solid-state, allows the rapid identification of RNA base-pair patterns.^[2,8,21,22]^ This information, even in the absence of site-specific assignments, provides accurate characterization of the RNA secondary structure, which is often sufficient to predict the function, the processing pathway and the stability of the RNA.^[23–25]^ Recently, ^1^H-detected ssNMR at 40 kHz MAS was used to observe inter-nucleotide Watson-Crick hydrogen bonds in the crystallized 23mer DIS-HIV-1. Unfortunately, the spectra lacked any site-specific resolution.^[26]^

Here we present two- and three-dimensional (2D and 3D) ^1^H-detected ssNMR experiments that allow rapid, nucleotide-specific detection of both Watson-Crick (G:C and A:U) and non-Watson-Crick (G:A and U:U) base pairs within hours (2D experiments) to days (3D experiments) at 100 kHz MAS. Furthermore, we describe ^1^H-detected ssNMR experiments that yield the complete resonance assignment of the nucleobases. This suite of solely ^1^H-detected ssNMR experiments is sufficient to accurately define the RNA secondary structure. Overall, the experimental approach described here paves the way to complete structure determination of RNA by ssNMR using only uniformly labelled samples.

## Results and Discussion

### Detection of inter-strand hydrogen bonds

In ssNMR Watson-Crick (WC) base pairs can be identified by the spatial proximity (2.6–2.9 Å) of the two nitrogen atoms involved in NH···N hydrogen bonds (U N3···N1 A; G N1···N3 C), using ^15^N-^15^N RFDR (radio-frequency-driven recoupling) correlations.^[8,21]^ However, this strategy is not suited to observing several non-Watson-Crick base pairs (e.g. 2-carbonyl-N3, 4-carbonyl-N3 U:U and trans-Hoogsteen/sugar-edge G:A^[27]^ base pairs), which either do not feature NH···N hydrogen bonds (U:U) or have longer distances between the donor and acceptor nitrogen atoms (G:A). More conveniently, NHHN and NHHC^[8,22,28]^ experiments correlate ^15^N/^13^C spins via magnetization transfer through their bound hydrogen atoms, utilizing proton spin diffusion (PSD) between hydrogen atoms that are close in space. In essence, these experiments rely on through-space ^1^H-^1^H correlations, similar to NOESY in solution-state NMR.^[29]^ Because at fast MAS rates (≥40 kHz) spin diffusion-based recoupling schemes are inefficient, here we instead used the RFDR scheme to exploit ^1^H-^1^H through-space correlations and measure cross-strand hydrogen bonds at 100 kHz MAS rate.^[30,31]^

We developed two different 3D ^1^H,^1^H RFDR correlation experiments, that allow the identification of Watson-Crick (G:C, A:U) and non-Watson-Crick (G:A, U:U) base pairs and also facilitate nucleobases assignment. The 2D versions of these experiments (**2D (H)N(HHC)H** and **2D (H)N(HHN)H**) at 100 kHz MAS are similar to the NHHC and NHHN^[8,22,28]^ experiments at low MAS rates (≤20 kHz), respectively, but the high sensitivity of proton detection allowed reduced experiment times of 12–15-hours, compared to the 60-hours experiment time needed for each of the NHHN/NHHC experiments.^[32]^ Moreover, the proton spectral dimension yields improved separation of the correlations in the resulting spectra. In these experiments, the magnetization is first transferred using a short CP period from 1H to 15N, where it evolves during t1, before being transferred back to ^1^H. ^1^H-^1^H inter-strand transfer across the base pair is then achieved by 0.48-ms (2D (H)N(HHN)H) or 0.4-ms (2D (H)N(HHC)H) ^1^H-^1^H RFDR mixing. The magnetization is then transferred to ^15^N (2D (H)N(HHN)H) or ^13^C (2D (H)N(HHC)H) by CP, on which the frequency of the heteronucleus can be encoded during *t*_*2*_ in the 3D version of the experiments. A final CP transfer brings the magnetization back to ^1^H for direct detection (Fig. 1c-f and Fig. S1a-b).

Both WC A:U and non-Watson-Crick G:A (trans-Hoogsteen/sugar-edge^[27]^) base pairs can be detected with the 2D (H)N(HHC)H experiment. In trans-Hoogsteen/sugar-edge G:A base pairs, the G-H1’ is close to the A-H6, leading to the corresponding A-N6–G-H1’ peak (Fig. 1c and g). In A:U base pairs, the proximity of A-H2 to the U-H3 (~2.8 Å) allows their straightforward identification through the U-N3–A-H2 correlations (Fig. 1d and h). As expected from the previously determined secondary structure,^[8]^ we observed two G:A and one A:U base pair in the 26mer box C/D RNA (Fig. 1g and h).

In G:C base pairs, the proximity of G-H1 to the C-H4 protons (~2.3–2.7 Å) allows for efficient ^1^H-^1^H magnetization transfer (Fig. 1e). Using the 2D (H)N(HHN)H experiment, we observed three out of the expected four G:C base pairs, identified by their C-N4–G-H1 and G-N1–C-H41/H42 cross peaks (Fig. 1i). In this experiment, the close proximity of intra-residue G-H1 and G-H2 protons (~2.1–2.3 Å) also results in intra-residue G-N1–H2 and G-N2–H1 correlations. While inter-strand C-N4–G-H1 correlations are well separated from the intra-residue G-N2–H1 correlations, the region of inter-strand G-N1–C-H41/H42 cross peaks overlaps with that of the intra-residue G-N1–H2 peaks, due to similar C-H41/H42 and G-H2 chemical shifts (Fig. 1i).

The relatively short distance between the two U-H3 imino protons in U:U base pairs (~2.9 Å) should allow their detection in the (H)N(HHN)H experiment (Fig. 1f). However, the presence of the predicted U3:U23 2-carbonyl-N3, 4-carbonyl-N3 base pair^[8]^ could not be confirmed in the 2D spectrum, due to the near-identical chemical shifts of U3-N3 and U23-N3. The presence of this base pair was confirmed later (Fig. 5e) in a 3D version of this experiment, which featured the evolution of both U-H3 proton frequencies.

In conclusion, these two 2D experiments allowed the rapid and almost complete identification of the base-pairing pattern in the 26mer box C/D RNA.

### Assignment of nucleobases

After initial assessment of the secondary structure of the 26mer box C/D RNA, we proceeded to the site-specific assignment of nucleobase spins, and thus base pairs, by ^1^H-detected ssNMR. Figure 2 shows 2D dipolar-based ^1^H-^13^C (Fig. 2a) and ^1^H-^15^N (Fig. 2b) cross polarization (CP)-HSQC spectra recorded for the nucleobases. With a few modifications, the experiment follows the scheme proposed by Rienstra and co-workers.^[33–35]^

**Figure 2.**
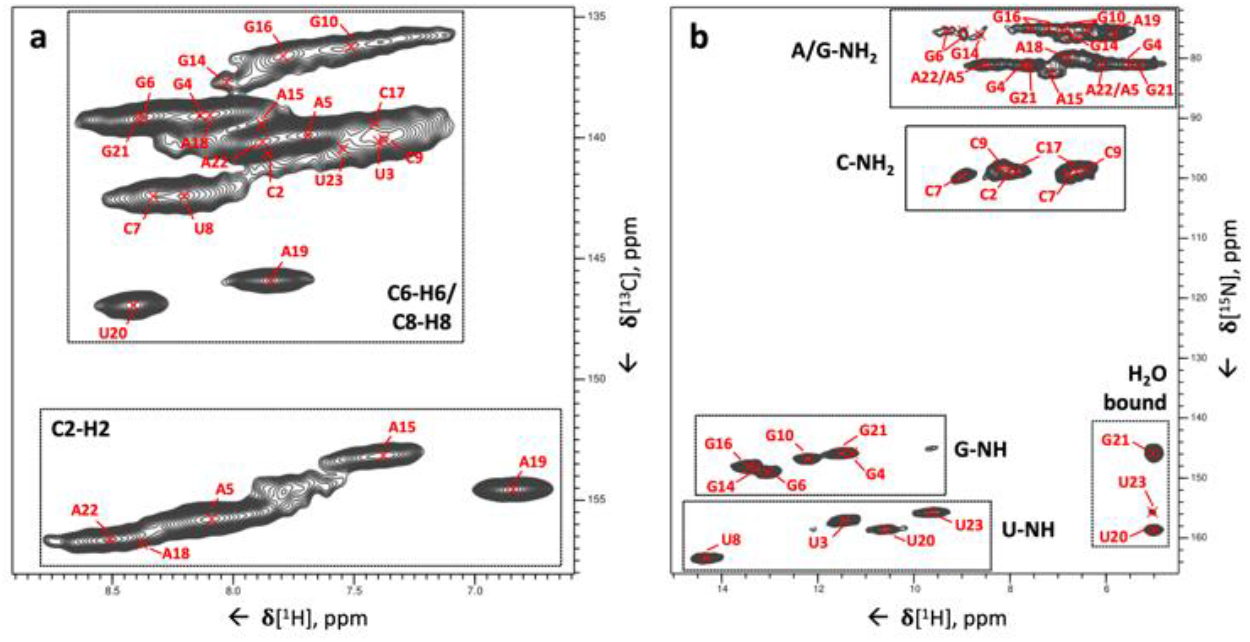
Two-dimensional ^1^H-^13^C and ^1^H-^15^N correlations of the 26mer box C/D RNA. (**a**) ^1^H-^13^C CP-HSQC spectrum tailored for nucleobase resonances showing C2–H2, C6–H6 and C8–H8 cross-peaks. The site-specific assignment of C8–H8 and C6–H6 cross-peaks is from our previous work,^[14]^ while that of the C2–H2 peaks has been obtained in this study. (**b**) ^1^H-^15^N CP-HSQC spectrum. The assignments have been obtained through the experiments developed here. The pulse sequence and phase cycle of the ^1^H-^15^N CP-HSQC experiment are given in Fig. S1c.

Our strategy for the complete assignment of RNA resonances in individual nucleotides by ^1^H detected ssNMR on uniformly ^13^C,^15^N-labelled RNA comprises of five steps. First, C1’-H1’ resonances are correlated with C6-H6 and C8-H8 resonances in pyrimidines and purines, respectively. Second ribose C-H resonances (C2’-H2’, C3’-H3’, C4’-H4’, C5’-H5’and -H5’’) are correlated to the C1’-H1’ resonances. These two steps were described previously^[14]^ and provided the assignment of 75% of the ribose spin systems as well as their connection to the pyrimidines C6-H6 and purines C8-H8. Third, all nucleobase carbon resonances are correlated to the previously assigned C6-H6 (pyrimidines) or C8-H8 (purines) groups. In the fourth step, the ^15^N-^1^H imino and amino resonances are correlated to the assigned carbons and C-H groups. Finally, the resonances of the non-protonated nitrogen atoms (N1, N3, N7 in purines and N3 in cytosine) are correlated to the assigned C-H groups. Here, we present experiments to accomplish steps 3–5 as well as to assign cross-strand hydrogen bonds. In the following spectra, for clarity, we annotate the spin systems according to their site-specific assignment, which we accomplished in previous work.^[7,8,14]^ The sequential assignment of RNA relies on through-space ^1^H-^1^H magnetization transfer, as explained previously^[8]^: experiments exploiting ^1^H-detection to accomplish sequential assignment of RNA are currently in development in our laboratory. The experiments presented here are complementary, allowing the complete assignment of nucleobase spin systems and the identification of the base pair pattern with spin-system resolution.

The **3D (H)CCH** experiment (Fig. S1d) accomplishes step 3. The experiment starts with a long-range ^1^H-^13^C CP transfer followed by evolution of ^13^C magnetization during *t*_*1*_. A selective ^13^C REBURP pulse^[36]^ with a bandwidth of ~50 ppm centered at ω(^13^C) = 160 ppm allows phase-cycled cancellation of the signals of the ribose ring, whose carbon spins are not affected by the pulse. The following 8 ms-long ^13^C-^13^C RFDR mixing period transfers ^13^C magnetization to nearby ^13^C atoms, whose chemical shift is recorded during *t*_*2*_. We used the ^13^C-^13^C RFDR mixing scheme^[30,31]^ to transfer magnetization among the nucleobase carbon spins (C2, C4, C5, C6, C8) because PDSD^[37]^ (proton- driven spin-diffusion) and DARR^[38]^ (dipolar-assisted rotational resonance) ^13^C-^13^C transfer schemes fail at high spinning rates. Finally, the ^13^C magnetization is transferred to the directly attached 1H via a second short-range CP and detected during *t*_*3*_ (Fig. 3a and j). This **3D (H)CCH** experiment correlates all carbons of the nucleobase to either the C8-H8 and C2-H2 groups in purines, or the C6-H6 and C5-H5 groups in pyrimidines. For instance, for the 26mer box C/D RNA, the experiment provided unambiguous assignment of all five adenosine C2-H2 resonances in the well-structured region (nucleotides 2–9 and 14–24) (Fig. 2a).

**Figure 3.**
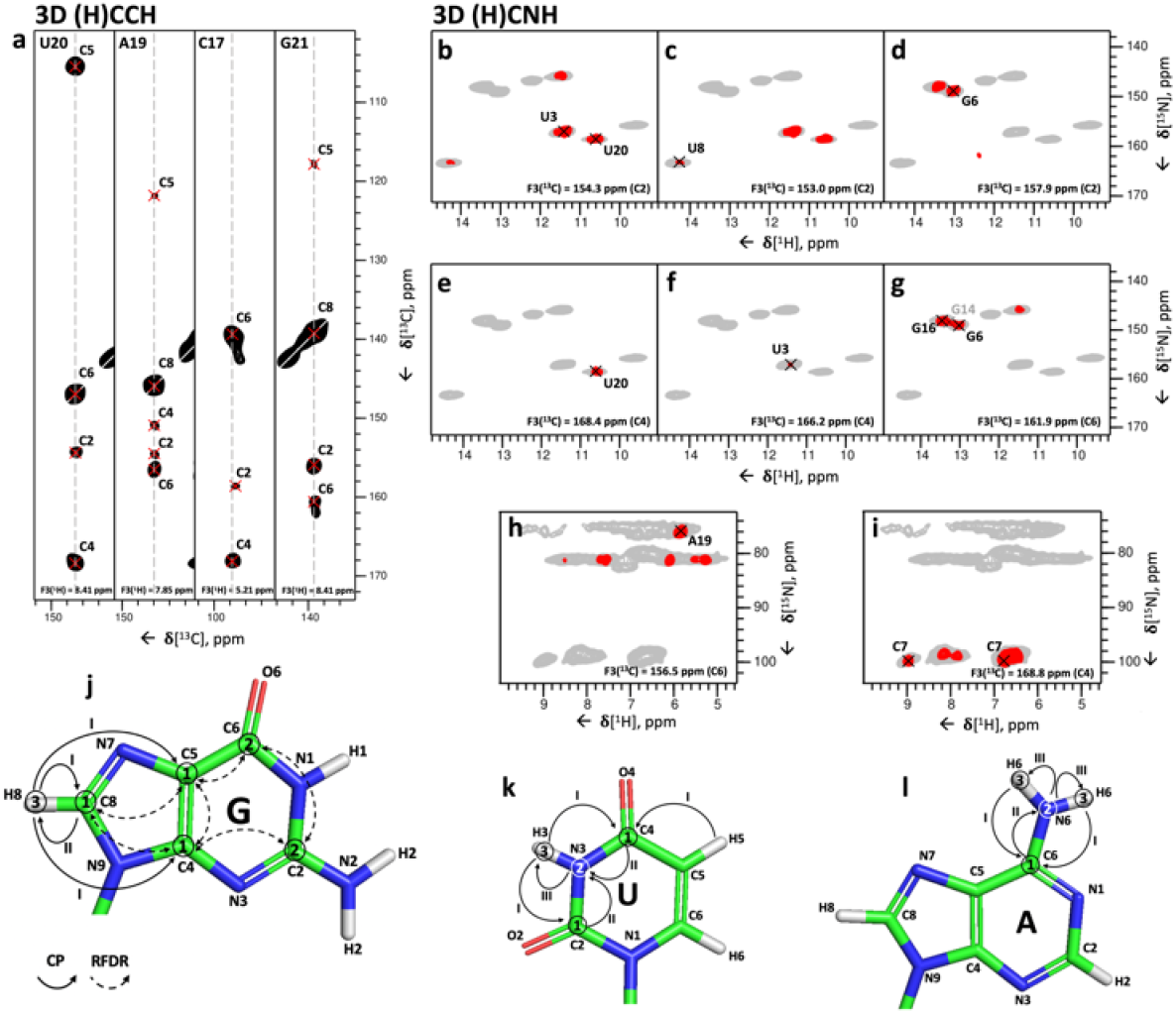
3D (H)CCH and 3D (H)CNH experiments for the correlation of nucleobase carbon, imino and amino resonances. (**a**) Representative 2D ^13^C-^13^C planes extracted from the 3D (H)CCH spectrum at the ^1^H frequencies of U20-H6, A19-H8, C17-H5 and G21-H8. Cross-peaks between U20-C6, A19-C8, C17-C5 and G2-C8 and all other respective base carbons can be identified. The experimental time was 127 hours. The pulse sequence and the phase cycle are given in Fig. S1d. (**b-i**) Representative 2D ^1^H-^15^N planes extracted from the imino-(b-g) or amino-(h-i) selective 3D(H)CNH spectra showing correlations of imino and amino nitrogen with carbon resonances in the nucleobase. The correlations shown are: U3/U20-C2–N3–H3 (**b**), U8-C2–N3–H3 (**c**), G6-C2–N1–H1 (**d**), U20-C4–N3– H3 (**e**), U3-C4–N3–H3 (**f**), G6-C6–N1–H1 (**g**), A19-C6–N6–H6 (**h**) and C7-C4–N4–H4 (**i**). In (g), the G14 peak is labelled in grey to indicated that the peak maximum is not in the plane shown here. The pulse sequence and the phase cycle of the experiment are given in Fig. S1e. For reference, in panels (b-i) the red contours of the 3D(H)CNH spectra are overlaid on the 2D ^1^H-^15^N CP-HSQC spectrum (in grey). (**j-l**) Magnetization transfer schemes of the 3D (H)CCH experiment shown for guanosine (G) (**j**) and of the 3D (H)CNH experiment shown for uridine (U) (**k**) and adenosine (A) (**l**). Encircled numbers indicate the chemical-shift evolution times (*t*_*1*_–*t*_*3*_) corresponding to the three spectral dimensions; roman numerals indicate CP transfer periods.

The fourth step of the assignment protocol is accomplished by the **3D (H)CNH** experiment (Fig. S1e). The experiment starts with a long-range ^1^H-^13^C CP transfer of 3 ms to transfer the ^1^H magnetization to non-protonated carbon atoms, whose chemical shifts are recorded during *t*_*1*_. Next, a 7-ms 13C-15N CP transfers the 13C magnetization to nitrogen atoms, whose chemical shifts are recorded in *t*_*2*_. Here, a selective 15N refocusing pulse (bandwidth of ~90 ppm), centered at either ω(^15^N) = 154 ppm or ω(^15^N) = 80 ppm after the *t*_*2*_ period selects for either imino (G-N1, U-N3) or amino (A-N6, G-N2, C-N4) nitrogen magnetization, respectively. Finally, the magnetization is transferred by CP from the nitrogen to the directly bound hydrogen for detection (Fig. 3k-l). The experiment yielded imino and amino-selective CX–NX–HX correlations in only 35 and 60 hours, respectively.

The combination of the 3D (H)CCH and the two 3D (H)CNH experiments allows assignment of the imino and amino resonances to individual nucleotides, provided that the chemical shift of at least one of the correlated carbons is unique to that nucleotide. This condition is not always fulfilled, due to the narrow chemical shift dispersion of A-C6, C-C4 and G-C2, which may hinder the nucleotide-specific assignment of A, C and G amino groups. Nevertheless, the imino-selective (H)CNH spectrum provided four U-C2/C4–N3–H3 and six G-C2/C6–N1–H1 correlations (Fig. 3b-g), allowing the unambiguous assignment of all four uridine imino resonances and six of the guanosine imino resonances present in the ^1^H-^15^N CP-HSQC spectrum (Fig. 2b, the peak of the nucleotide G24 is missing). In the amino-selective (H)CNH spectrum, we detected four sets of cytidine C4–N4–H4 correlations, but the low chemical shift dispersion of the carbon C4 resonances hindered the assignment of the amino groups of the C2, C9 and C17 nucleotides (Fig. 4a), and only the amino group of C7 could be assigned unambiguously (Fig. 3i). Similarly, due to the low chemical shift dispersion of A-C6 and G-C2 spins, the adenosine and guanosine amino resonances were poorly resolved. The 3D (H)CNH spectrum yielded the assignment of the amino resonance of nucleotide A19 only, which has a unique carbon C6 chemical shift (156.5 ppm, Fig. 3h). The unambiguous assignment of the other adenosine amino groups was hindered by almost identical C6 chemical shifts, such as those of A15 and A18 (~157.9–158.0 ppm, Fig. 4b) or A5 and A22 (~157.3–157.4 ppm, Fig. 4c).

**Figure 4.**
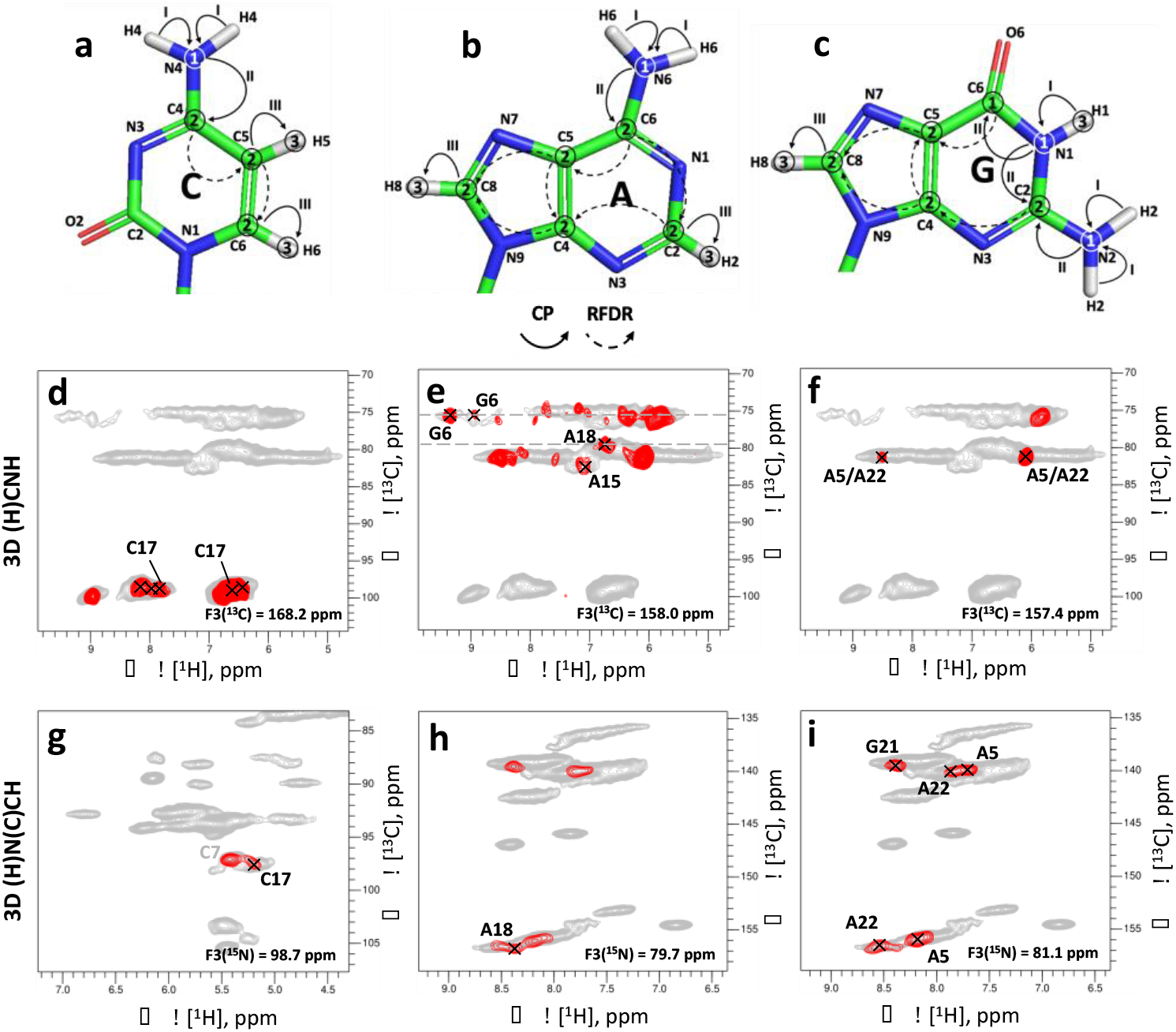
3D (H)N(C)CH experiment for assignment of nucleobase amino resonances. (**a-c**) Magnetization transfer schemes of the 3D (H)N(C)CH experiment shown for cytidine (**a**) adenosine (**b**) and guanosine (**c**). Encircled numbers indicate the chemical-shift evolution times (*t*_*1*_–*t*_*3*_) corresponding to the three spectral dimensions; roman numerals indicate CP transfer periods. (**d-f**) Representative 2D ^1^H-^15^N planes extracted from the amino-selective 3D (H)CNH experiment (introduced in Fig. 3) showing cytidine C4–N4–H4 (**d**), adenosine C6–N6–H6 and guanosine C2– N2–H2 (**e**) and adenosine C6–N6–H6 (**f**) correlations, which could not be assigned unambiguously in this experiment. (**g-i**) Representative 2D ^1^H-^13^C planes extracted from the 3D (H)N(C)CH spectrum showing cytidine N4–C5–H5 (**g**), adenosine N6–C2–H2 and N6–C8–H8 and guanosine N2–C8–H8 (**h** and **i**) correlations, which provide the assignments that could not be resolved from the 3D (H)CNH experiment alone. In (g), the C7 peak is labelled in grey to indicated that the peak maximum is not in the plane shown here. The pulse program and the phase cycle of the 3D (H)N(C)CH experiment are given in Fig. S1f. For reference, in panels (d-f) the red contours of the 3D (H)CNH spectrum are overlaid on the 2D ^1^H-^15^N CP-HSQC spectrum (in grey), and in panels (g-i) the red contours of the 3D (H)N(C)CH spectrum are overlaid on 2D ^1^H-^13^C CP-HSQC spectra (in grey) tailored either for the ribose/C5-H5 (g) or the C2-H2/C6-H6/C8-H8 spectral regions (h-i).

To overcome the problems imposed by the limited chemical shift dispersion described above, we developed **3D (H)N(C)CH** and **H(NC)CH** experiments, with the aim of exploiting the better resolution of protonated carbons to resolve either the nitrogen or the hydrogen chemical shifts of imino and amino groups. The 3D (H)N(C)CH experiment (Fig. S1f) starts with a ^1^H-^15^N CP transfer of 1 ms, followed by evolution of ^15^N chemical shifts during *t*_*1*_. The following 10 ms-long 13C-15N CP period transfers magnetization from the nitrogen to the directly attached carbon. The selective REBURP refocusing pulse (bandwidth of ~50 ppm) centered at ω(^13^C) = 160 ppm selects for nucleobase carbon magnetization, which is then transferred to all adjacent carbons, including the protonated ones, via a 14 ms-long ^13^C-^13^C RFDR mixing period. Finally, after evolution of the carbon chemical shift during *t*_*2*_, the magnetization is transferred from the protonated carbon atoms to the directly attached hydrogen atoms for detection (Fig. 4a-c). In the (H)N(C)CH experiment, the frequencies of the imino and amino nitrogen atoms are correlated to the C5-H5 or C6-H6 groups in pyrimidines and to the C2-H2 or C8-H8 groups in purines (Fig. 4g-i). The poor resolution of the C8-H8 peaks of guanosines makes the assignment of their imino and amino nitrogen atoms more difficult than for the other nucleobases, where the assignment is facilitated by the well-resolved C2-H2 (adenosines) and C5-H5 (cytidine and uridine) groups.

Although this experiment has relatively low sensitivity, due to the long and inefficient ^15^N-^13^C transfer, correlations of imino nitrogen atoms were observed for four uridines (U3-, U8-, U20-, U23-N3 to C5-H5 groups and U3-, U20-N3 to C6-H6 groups) and three guanosines (G4-, G6-, G21-N1 to C8-H8 groups), whereas correlations of amino nitrogen atoms were obtained for two cytidines (C7-, C17-N4 to C5-H5 and C6-H6 groups), four adenosines (A5-, A18-, A19-, A22-N6 to C2-H2 groups and A5-, A19-, A22-N6 to C8-H8 groups) and three guanosines (G4-, G6-, G21-N2 to C8-H8 groups) after a measurement time of 80 hours. These correlations allowed us to resolve the ambiguity in the assignment of A15 and A18: while in the (H)CNH experiment, the two corresponding peaks at N6-H6 chemical shifts of 82.6/7.1 ppm and 79.7/6.8 ppm cannot be attributed to one or other of A15 and A18 due to the degeneracy in the C6 chemical shift (Fig. 4e), the presence of a cross-peak between the C2-H2 of A18 with an N6 at 79.7 (Fig. 4h) unambiguously attributes the N6-H6 group at 79.7/6.8 ppm to A18. Furthermore, the presence of cross peaks between an N6-H6 group with a nitrogen chemical shift at 81.1 ppm and both C8-H8 and C2-H2 groups of both A5 and A22 revealed that the amino groups of A5 and A22 overlap (Fig. 4f and i). Finally, we observed N4–C5–H5 cross-peaks for C7 and C17 (Fig. 4g); however, we find another experiment to be better suited for the assignment of cytidine amino groups (*vide infra*). By contrast, the sensitivity of the 3D H(N)(C)CH experiment, where the chemical shifts of ^1^H instead of ^15^N were evolved during *t*_*1*_, remained very low after 62 hours of measurement time. This can be explained by the short coherence lifetimes of the imino and amino protons (imino protons *T*_*2*_’ ≈ 4 ms, amino protons *T*_*2*_’≈ 1 ms at 100 kHz MAS).

For the assignment of the cytidine amino groups, we used the **3D (H)N(HH)CH** experiment (Fig. S1a), which utilizes the efficient ^1^H-^1^H transfer between the proximal H5 and H41/H42 hydrogens (~2.4–2.7 Å) and provides the same N4–C5–H5 correlations as the 3D (H)N(C)CH experiment (Fig. S2a). The efficiency of the ^1^H-^1^H RFDR mixing (0.48 ms-long) was superior to that of the ^13^C-^13^C RFDR mixing, leading to a higher sensitivity of the 3D (H)N(HH)CH experiment compared to the 3D (H)N(C)CH experiment. The 3D (H)N(HH)CH experiment provided N4–C5–H5 correlations of the four cytidines C2, C7, C9, C17 (excluding C26 which is not visible in any ssNMR spectrum, Fig. S2c-f) and allowed unambiguous assignment of the amino resonances of nucleotides C2, C9 and C17 (Fig. S2d-f).

The 3D (H)N(C)CH experiment did not show any N2–C8–H8 cross-peaks for G10, G14, G16 and G24. To assign the guanosine amino groups, we exploited the intra-residue proximity of the H1 and H2 hydrogen atoms (~2.1–2.3 Å) in a **3D (H)N(HH)NH** experiment (Fig. S1b and Fig. S2b), which utilizes the same magnetization transfer pathway as the 2D (H)N(HHN)H experiment (Fig. 1i) but features an additional ^15^N evolution time before the final magnetization transfer to ^1^H for direct detection, thereby yielding N2–N1–H1 and N1–N2–H2 cross-peaks. In the 3D (H)N(HH)NH spectrum we observed N2–N1–H1 cross-peaks for the nucleotides G4, G6, G10, G14, G16 and G21 (Fig. S2g-j). Combining the information on N2 and C2 chemical shifts obtained from the 3D (H)N(HH)NH and amino-selective (H)CNH experiments, we were able to unambiguously assign guanosine amino resonances for five out of the six guanosines in the structured region of the 26mer box C/D RNA (nucleotides 2–9 and 14–24) and of G10 in the apical loop (Fig. S2g-j). The N-H resonances of G24 at the open end of the helix were not detectable in any of the spectra.

In total, using the combination of experiments presented in Fig. 3–4 and S2 we could assign all imino groups of the 19 structured nucleotides (2–10 and 14–24) with only G24 missing, as well as the amino groups of all cytidines, five out of six guanosines (excluding G24 in the helix end) and all adenosines.

In the final step, we assigned the remaining nitrogen resonances (N1, N3, N7 in purines) using a modified version of the base-tuned **3D (H)NCH** experiment developed previously.^[14]^ We shifted the ^15^N carrier position from 160 to 190 ppm to allow effective manipulation of the purine N1, N3 and N7 resonances. In addition, after *t*_*2*_, we applied a selective ^13^C refocusing pulse (bandwidth of 50 ppm) centered at ω(^13^C) = 160 ppm to select for nucleobase ^13^C resonances. The experiment yields N7–C8–H8 and N9–C8–H8 correlations in purines, N1–C2–H2 and N3–C2–H2 correlations in adenosines and N1–C6–H6 correlations in pyrimidines (Fig. S1g and Fig. S3) and provided the assignment of all missing nitrogen atoms in purines in 56 hours of measurement time. Finally, to assign N3 resonances in cytidines involved in base pairs, we recorded a 2D ^1^H-^15^N-HSQC experiments with long-range ^1^H-^15^N/^15^N-^1^H CP transfers of 8 ms, which allows magnetization transfer from guanosines H1s to cytidine N3s involved in WC base pairs (Fig. S1c and Fig. S4). However, in this experiment C-N3–G-H1 correlations could be obtained only for the G6:C17 base pair (Fig. S4b).

### Assignment of base pairs

The two-dimensional spectra of Fig. 1 allow the assignment of base pairs to specific pairs of spin-systems. The WC A:U base pair (Fig. 1h) can be assigned to the uridine spin-system with U-N3 = 163.3 ppm (U8) and to the adenosine spin-system with A-H2 = 7.4 ppm (A15). Similarly, the three WC G:C base pairs belong to three pairs of spin-systems with G-H1 = 13.0 ppm and C-N4 = 98.8 ppm (G6:C17), G-H1 = 13.3 ppm and C-N4 = 98.2 ppm (G14:C9), G-H1 = 13.4 ppm and C-N4 = 99.4 ppm (G16:C7). Finally, the two A-N6–G-H1’ correlations specific for trans-Hoogsteen/sugar-edge G:A base pairs (Fig. 1g) involve one guanosine spin-system with the unique H1’ chemical shift of 6.1 ppm (G21) and one other guanosine spin-system with the non-unique chemical shift of 5.7 ppm (G4 or G10).^[14]^ The identity of the adenosine spin-system remains ambiguous as the two adenosine spin-systems involved in the G:A base pairs have the almost identical N6 chemical shifts of 81.1–81.2 ppm (A5 and A22).

To be able to assign the base pairs to individual spin-systems in larger RNAs with more spectral crowding, we tested 3D versions of the experiments of Fig. 1. The 3D (H)N(HH)CH experiment, with an additional evolution time for ^13^C frequencies (as in Fig. S2c–f), was acquired with a ^1^H-^1^H RFDR mixing time of 0.96 ms (compared to that of 0.48 ms used in the 2D spectrum of Fig. 1). One U-N3–A-C2–H2 cross-peak confirmed the U8:A15 base pair (Fig. 5a). In addition, the different C1’chemical shifts of G4 and G10 allowed identification of G4 as the nucleotide involved in the second G:A base pair. Using the ^1^H-^1^H RFDR mixing time of 0.96 ms, we observed additional cross-peaks between nitrogen atoms corresponding to either G-N2 or A-N6 and the C8-H8 groups of two adenosines at 139.9/7.7 ppm and 140.1/7.9 ppm (A5 and A22) (Fig. 5c). If these cross-peaks originated from G-N2, they would be indicative of G:A base pairs; alternatively, they could originate from A-N6 through intra-residue magnetization transfer. To verify the nature of these cross-peaks we evaluated intra-residue and inter-strand A-H6–H8 and G-H2–A-H8 distances in the ssNMR-derived structure of this RNA^[15]^ as well as in a few other known RNA structures, that contain trans-Hoogsteen/sugar-edge G:A base pairs^[39,40]^ (Table S4). Inter-strand G-H2–A-H8 distances (~ 3.1 Å) are significantly shorter than intra-residue A-H6–H8 distances (~ 5.0 Å); furthermore, we did not observe any intra-residue A-H6–H8 correlations for any other adenosine spin system apart from A5 and A22. Thus, we concluded that the correlations observed in the 3D (H)N(HH)CH spectrum are specific for trans-Hoogsteen/sugar-edge G:A base pairs. The longer distance of inter-strand G-H2–A-H8 hydrogens compared to that of inter-strand G-H1’–A-H6 hydrogens (~ 2.4 Å) explains why the G-H2–A-H8 cross-peaks were observable only when using the longer ^1^H-^1^H RFDR mixing time of 0.96 ms.

**Figure 5.**
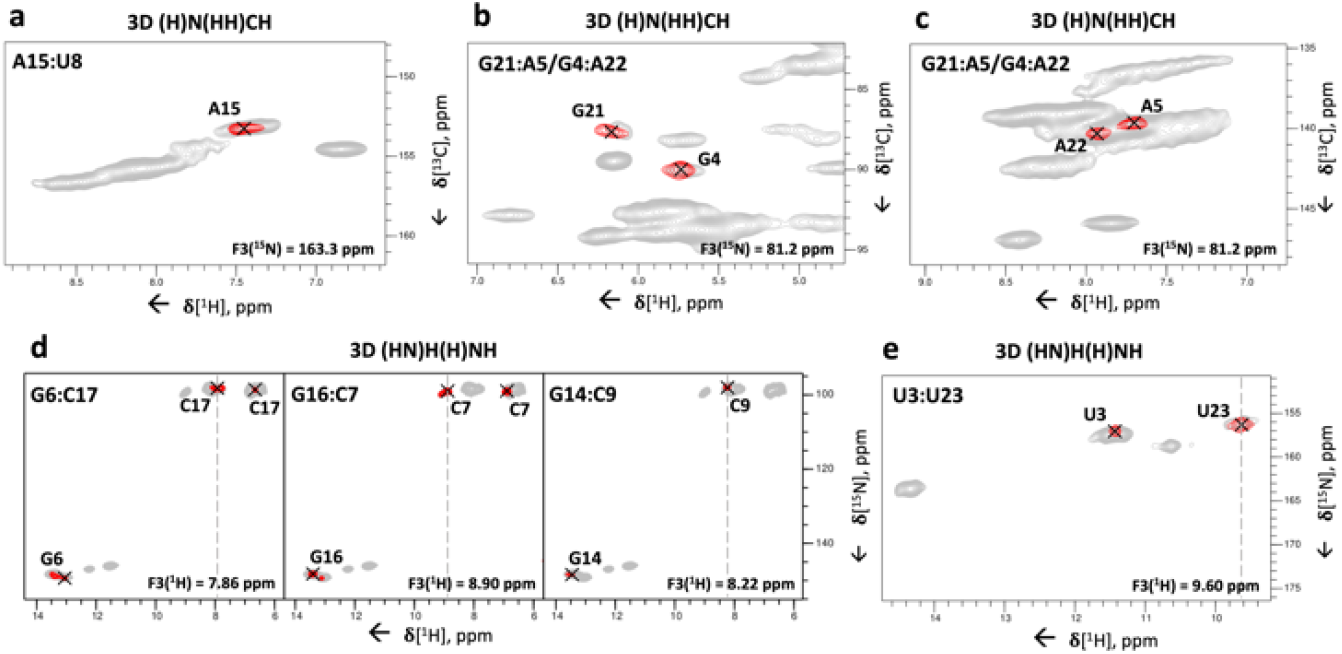
3D correlation experiments for the assignment of base pairs to individual spin systems. (**a-c**) Representative 2D ^1^H-^13^C planes extracted from the 3D (H)N(HH)CH spectrum to identify nucleotides involved in base pairs. **(a)** U-N3–A-C2–H2 cross-peak of the base pair U8:A15; **(b)** A-N6–G-C1’–H1’ and (**c**) G-N2–A-C8–H8 cross-peaks of the base pairs G4:A22 and A5:G21. The ^1^H-^1^H RFDR mixing time was 0.96 ms. (**d-e**) Representative 2D ^1^H-^15^N planes extracted from the 3D (HN)H(H)NH spectrum showing the inter-strand C-H41/H42–G-N1–H1 and intra-residue C-H41–N4–H42 cross-peaks for the base pairs G6:C17, C7:G16 and C9:G14 (**d**); the inter-strand U-H3–U-N3–H3 cross-peak of the base pair U3:U23 (**e**). The ^1^H-^1^H RFDR mixing time was 0.48 ms. For reference, in panels (a-c) the red contours of the 3D (H)N(HH)CH spectrum are overlaid onto the 2D ^1^H-^13^C CP-HSQC spectra (in grey) tailored either for the base (a) or ribose/C5-H5 spectral regions (b-c). In panels (d-e), the red contours of the 3D (HN)H(H)NH spectrum are overlaid onto the 2D ^15^N-^1^H CP-HSQC spectrum (in grey). The dashed lines indicate the positions of the diagonal peaks in the ^1^H-^1^H plane. Pulse sequences and phase cycles are given in Fig. S1a and b.

We also implemented two different 3D versions of the (H)N(HHN)H experiment (Fig. S1b), where we utilized a ^1^H-^1^H RFDR mixing time of 0.48 ms and measured either ^1^H, ^15^N, ^1^H or ^15^N, ^15^N, ^1^H frequencies (as in Fig. S2g–j) and. Both experiments showed good sensitivity (Fig. 5d and Fig. S5), but the **3D (HN)H(H)NH** experiment, with acquisition of ^1^H, ^15^N, ^1^H frequencies, had better resolution of the base pairs. C-H41/H42– G-N1–H1 and C-N4–G-N1–H1 cross-peaks were obtained for three out of the four G:C base pairs of the 26mer Box C/D RNA (Fig. 5d and Fig. S5). No cross-peak was observed for the terminal base pair C2:G24 due to either poor signal-to-noise or to fraying of the helix. The 3D (HN)H(H)NH experiment delivered also intra-residue C-H41–N4–H42 cross-peaks, which correlated directly the H41 and H42 resonances.

Finally, the 3D (HN)H(H)NH experiment yielded the assignment of the U:U base pair by separating the resonances of the degenerate N3 atoms via the different chemical shifts of the H3 hydrogens: we observed a cross-peak between two uridine imino protons at 9.6 and 11.4 ppm, which confirmed the presence of the predicted U:U base pair^[8]^ between the two unique uridine spin-systems containing these imino protons (U3:U23, Fig. 5e).

## Conclusion

In summary, we have established a set of experiments that accomplishes the rapid identification of RNA base pair patterns, the complete assignment of all nucleobase spin-systems (Table S5) and the assignment of the base pairs to individual spin-systems. The experimental workflow requires a single uniformly-labelled RNA sample and uses ^1^H-detected MAS ssNMR at ultrafast (≥100 kHz) spinning rates. We are currently developing strategies for the determination of structural restraints based solely on ^1^H-detected ssNMR applied to a single uniformly-labelled RNA sample. These developments demonstrate that ^1^H-detection has the potential to significantly strengthen the role of ssNMR in RNA structural biology and will pave the way for the determination of RNA structures within large biomolecular complexes.

## Acknowledgements

This work was supported by the Deutsche Forschungsgemeinschaft (DFG grant CA294/21-1 to TC).

