## Supporting information for "Identification of RNA base pairs and complete assignment of nucleobase resonances by proton-detected solid-state NMR spectroscopy at 100 kHz MAS"

**Abstract:** Knowledge of RNA structure, either in isolation or in complex, is fundamental to understand the mechanism of cellular processes. Solid-state NMR (ssNMR) is applicable to high molecular-weight complexes and does not require crystallization; thus, it is well-suited to study RNA as part of large multicomponent assemblies. Recently, we solved the first structures of both RNA and an RNA–protein complex by ssNMR using conventional $^{13}$C- and $^{15}$N-detection. This approach is limited by the severe overlap of the RNA peaks together with the low sensitivity of multidimensional experiments. Here, we overcome the limitations in sensitivity and resolution by using $^1$H-detection at fast MAS rates. We develop experiments that allow the identification of complete nucleobase spin-systems together with their site-specific base pair pattern using sub-milligram quantities of one uniformly labelled RNA sample. These experiments provide rapid access to RNA secondary structure by ssNMR in protein–RNA complexes of any size.

### Table of Contents


1. **Experimental Procedures** 2
   - **Protein expression and purification** 2
   - **RNA synthesis** 2
   - **Assembly of L7Ae–26mer box C/D RNA complex** 2
   - **NMR spectroscopy** 3
2. **Supplementary figures** 5
   - **Pulse sequences of all experiments** 5
   - **3D (H)N(HH)CH and 3D (H)N(HH)NH experiments for the assignment of nucleobase amino resonances** 6
   - **Assignment of non-protonated nucleobase nitrogen chemical shifts by the 3D (H)NCH experiment** 7
   - **Assignment of C-N3 resonances by the 2D long-range $^1$H-$^{15}$N CP-HSQC experiment** 8
   - **Detection of G:C base pairs by the 3D (H)N(HH)NH experiment** 9
3. **Supplementary tables** 10
   - **Acquisition parameters and evolution times of all spectra** 10
   - **Summary of cross-polarization conditions** 11
   - **Fourier processing parameters of all spectra** 13
   - **Inter-strand and intra-residue distances in trans-Hoogsteen/sugar-edge G:A base pairs** 14
   - **Isotropic chemical shifts of the 26mer box C/D RNA in the L7Ae–box C/D RNA complex** 15
4. **References** 15
5. **Author contributions** 16

### Experimental Procedures

**Protein expression and purification.** The L7Ae protein from *Pf* was expressed and purified through a denaturing purification protocol, as described in detail elsewhere.[1] Briefly, L7Ae was expressed from the pET-M11 expression vector, containing an N-terminal hexa-histidine tag, followed by a TEV (tobacco etch virus) protease cleavage site, in *Escherichia coli* BL21 (DE3) cells using LB medium.[2] Cells were grown at 37 °C and expression was induced at $OD_{600}$ = 0.6–0.8 by addition of 0.6 mM isopropyl-β-D-thiogalactopyranoside (IPTG). Expression was continued for 16–20 hours at 18 °C. Cells were harvested by centrifugation at 4000 g at 4 °C and lysed by sonication in the lysis buffer (50 mM Tris-HCl pH 7.5, 1 M NaCl, 2 mM β-mercaptoethanol (BME), 10 mM imidazole) with the addition of 0.2 mg/mL lysozyme, 40 μg/mL DNAse I and 1x protease inhibitor cocktail (Roche). The centrifuged (45 min, 4 °C, 32000 g) and filtered (0.45 μm) lysate was mixed with 8 M guanidinium chloride (GuHCl) pH 7.5 at a volume ratio of 1:3. The L7Ae protein was then purified by immobilized-metal-ion affinity chromatography (IMAC) with a HisTrap HP 5 mL column (GE Healthcare). The column was equilibrated with denaturing buffer (50 mM Tris-HCl, pH 7.5, 6 M GuHCl, 1 M NaCl, 2 mM BME, 10 mM imidazole); the protein solution was applied to the column and washed with 5 column volumes (CV) of denaturing buffer prior to protein refolding on the column by a 40 CV buffer gradient, which brought the column back into the lysis buffer. L7Ae was eluted with a 0–100 % gradient of elution buffer (50 mM Tris-HCl pH 7.5, 1 M NaCl, 2 mM BME, 300 mM imidazole) over 4 CVs. The tagged L7Ae protein was incubated with TEV protease (1:30 protease:protein ratio) overnight at room temperature in lysis buffer. The cleaved protein was isolated by reverse IMAC using a HisTrap HP column and further purified by size-exclusion chromatography (SEC) with a Superdex75 Increase 10/300 column (GE Healthcare) in SEC buffer (50 mM Tris-HCl pH 7.5, 1 M NaCl, 2 mM BME).

**RNA synthesis.** Uniformly $^{1}H,^{13}C,^{15}N$-labeled 26mer box C/D RNA (5´-GCUGAGCUCGAAAGAGCA-AUGAUGUC-3´) was prepared by *in vitro* transcription with T7 polymerase (produced in house), $^{13}C$, $^{15}N$ labeled NTPs and plasmid DNA template in transcription buffer (40 mM Tris-HCl pH = 8.0, 1 mM spermidine, 10 mM DTT, 0.01 % Triton X-100 and 0.1 % PEG8000).[3] Transcription yields were optimized in 20 μL small-scale reactions in micro-crystallization 96-well plates with varying concentrations of $MgCl_2$ (20 to 50 mM), rNTPs (10 to 40 mM), plasmid DNA (25 to 200 ng/μL) and *in-house* produced T7 polymerase (25 to 100 ng/μL).[3] $^{13}C$ and $^{15}N$ labeling was achieved using commercially available $^{13}C,^{15}N$-labeled rNTPs (Silantes). Small-scale reactions were incubated for 2 to 3 h at 37 °C and analyzed via analytical polyacrylamide gel electrophoresis (PAGE), using ethidiumbromid staining and UV visualization. The conditions leading to the best RNA yield were used for preparative-scale reactions. Thermostable inorganic pyrophosphatase (TIPP) (NEB) was added to the reaction to remove pyrophosphate. Preparative transcription reactions were stopped after 6 h by addition of EDTA (final concentration 50 mM). The RNA was purified by denaturing 15 % PAGE and extracted from the gel using the 'crush and soak' method in RNA elution buffer (40 mM MOPS pH 6.0, 11 mM EDTA) for 16 h at 4 °C prior to EtOH/NaCl precipitation. The pure RNA was dissolved in water.[1]

**Assembly of L7Ae–26mer box C/D RNA complex**. The individual components of the complex were tested for RNAse contamination using the RNAseAlert™ Lab Test Kit prior to complex assembly. The L7Ae–26mer box C/D RNA complex was assembled by mixing L7Ae protein and 26mer box C/D RNA in a 1.1:1 molar ratio. The mixture was incubated at 80 °C for 15 min, slowly cooled to room temperature and then purified by SEC on a Superdex75 Increase 10/300 column in RNP buffer (25 mM HEPES pH 7.5, 120 mM NaCl). The pure protein–RNA complex was concentrated to ~20 mg/ml in RNP buffer and subsequently mixed with an equal volume of precipitation solution (100 mM sodium acetate, 30 % PEG 400 in 100 mM HEPES, pH 7.5 in 100 % $H_2O$ or 50 % $D_2O$) as described elsewhere.[4–7] The sample was micro-crystallized by slow precipitation using a SpeedVac concentrator at room temperature for ~2 hours. The complex precipitated at approximately half the starting volume. The precipitate was packed into a 0.81-mm MAS ssNMR rotor by centrifugation at 80000 g for 2 h.

**NMR spectroscopy.** All NMR experiments were acquired on a Bruker Avance III HD NMR spectrometer operating at a $^{1}$H field-strength of 850 MHz and equipped with a 0.81-mm triple-resonance HCN probe developed in the Samoson laboratory (https://www.nmri.eu/).[8] All presented spectra were recorded with samples prepared in 100 % $H_2O$, except those shown in Fig. 6d and e, which were recorded on a sample prepared in 50 % $D_2O$. For observation of the $NH_2$ groups in the 3D (HN)H(H)NH spectrum, we chose to record the spectrum in a buffer containing 50 % $D_2O$, sacrificing 10–20 % peak intensity in favor of ~20–25 % narrower linewidths, resulting from longer proton coherence lifetimes. However, the quality of the spectrum did not significantly improve compared to the spectrum recorded in 100 % $H_2O$ (shown in Fig. S4).

The final temperature of the assignment experiments was optimized by inspecting 2D $^{1}$H-$^{15}$N HSQCs recorded at temperatures of 260 K, 265 K, 275 K, 280 K and 285 K. The $^{1}$H linewidths of a few nucleotides (e.g. G10, A15, A18, A19) became significantly broader at lower temperatures, while other nucleotides (e.g. U3, C7, U20) were unaffected by temperature changes. Conversely, the signal-to-noise ratio of most peaks was worse at higher temperatures (e.g. 280 K and 285 K). Thus, we chose the temperature of 275 K as a compromise between the temperature-dependent linewidths and sensitivity. All spectra for resonance assignment were recorded at a magic-angle spinning rate of 100 kHz.

WALTZ-16[9] decoupling with $\nu = \gamma B_1/2\pi = 10\,\mathrm{kHz}$ was used for $^{1}$H decoupling during indirect heteronuclear evolution periods (e.g. $^{13}$C, $^{15}$N). WALTZ-16 was also used for $^{15}$N decoupling during evolution of either $^{1}$H or $^{13}$C spins, including the direct acquisition time. DIPSI-3[10] decoupling with $\nu = \gamma B_1/2\pi = 20\,\mathrm{kHz}$ was used for $^{13}$C decoupling during either $^{1}$H or $^{15}$N evolution times. The MISSISSIPPI[11] scheme ($\nu = \gamma B_1/2\pi = 40\,\mathrm{kHz}$) was employed to suppress the residual $H_2O$ signal during z-storage of $^{13}$C or $^{15}$N magnetization.

The shapes, offsets, durations and powers employed for CP transfers are summarized in Table S2. Selective $^{13}$C and $^{15}$N refocusing pulses from the BURP family[12] were applied to eliminate signals outside the region of interest. The $^{15}$N and $^{13}$C carrier frequencies are summarized in Table S1.

$^{13}$C chemical shifts were referenced as described by Morcombe and Zilm.[13] $^{15}$N and $^{1}$H chemical shifts were referenced indirectly using chemical shift referencing ratios from the work of Markley *et al*.[14] Spectra were processed with NMRPipe[15] and visualized and assigned with CcpNmr Analysis.[16]

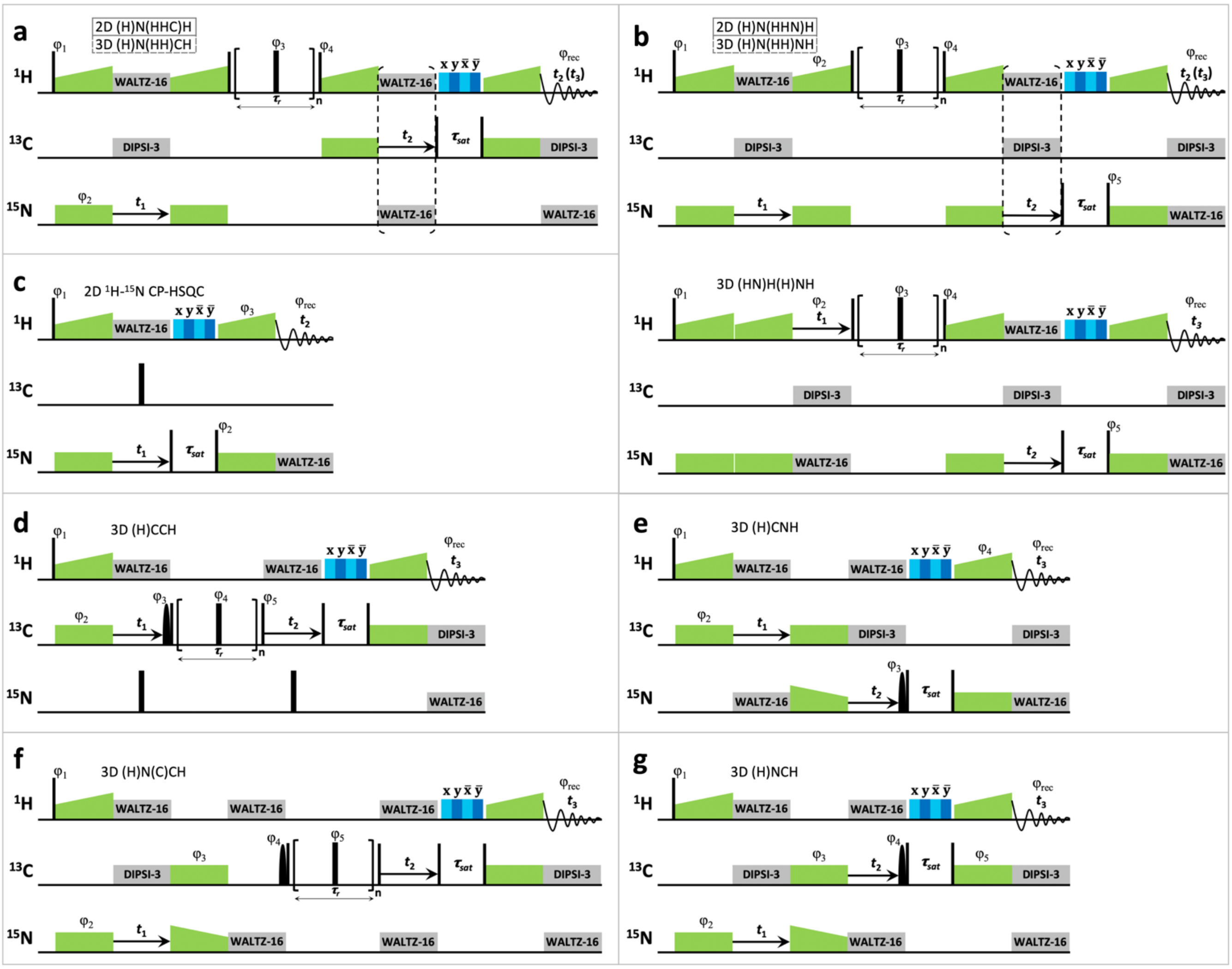


**Figure S1. Pulse sequences of all experiments.** (**a**) 2D (H)N(HHC)H and 3D (H)N(HH)CH experiments. In the 3D experiment the $^{13}$C chemical shifts were recorded during $t_2$. The phase cycle was $\varphi_1$ = y, –y; $\varphi_2$ = 2(x), 2(–x); $\varphi_3$ = x, y, x, y, y, x, y, x; $\varphi_4$ = 4(y), 4(–y); $\varphi_{rec}$ = x, –x, –x, x, –x, x, x, –x. (**b**) 2D (H)N(HHN)H, 3D (H)N(HH)NH and 3D (HN)H(H)NH experiments. In the 3D experiments the $^{15}$N chemical shifts were recorded during $t_2$. In $t_1$ either $^{15}$N (3D (H)N(HH)NH) or $^{1}$H (3D (HN)H(H)NH) chemical shifts were recorded. The phase cycle was $\varphi_1$ = y, –y; $\varphi_2$ = 2(y), 2(–y); $\varphi_3$ = x, y, x, y, y, x, y, x; $\varphi_4$ = 4(y), 4(–y); $\varphi_5$ = 8(y), 8(–y); $\varphi_{rec}$ = x, –x, –x, x, –x, x, x, –x, –x, x, x, –x, x, –x, –x, x. (**c**) 2D $^{1}$H-$^{15}$N CP-HSQC experiment. The phase cycle was $\varphi_1$ = y, –y; $\varphi_2$ = 2(x), 2(–x); $\varphi_3$ = 4(y), 4(–y) $\varphi_{rec}$ = y, –y, –y, y, –y, y, y, –y. (**d**) Base-selective 3D (H)CCH experiment. The phase cycle was $\varphi_1$ = 2(y), 2(–y); $\varphi_2$ = y, –y; $\varphi_3$ = 8(x), 8(y); $\varphi_4$ = x, y, x, y, y, x, y, x; $\varphi_5$ = 4(x), 4(–x); $\varphi_{rec}$ = x, –x, –x, x, –x, x, x, –x, –x, x, x, –x, x, –x, –x, x. (**e**) Amino- and imino-selective 3D (H)CNH experiment. The phase cycle was $\varphi_1$ = y, –y; $\varphi_2$ = 2(y), 2(–y); $\varphi_3$ = 4(x), 4(y), 4(–x), 4(–y); $\varphi_4$ = 16(y), 16(–y); $\varphi_{rec}$ = 2(–y, y, y, –y, y, –y, –y, y), 2(y, –y, –y, y, –y, y, y, –y). (**f**) Base-selective 3D (H)N(C)CH experiment. The phase cycle was $\varphi_1$ = 2(y), 2(–y); $\varphi_2$ = y, –y; $\varphi_3$ = 2(x), 2(–x); $\varphi_4$ = 4(x), 4(y); $\varphi_5$ = x, y, x, y, y, x, y, x; $\varphi_{rec}$ = x, –x, –x, x, –x, x, x, –x. (**g**) Base-selective (H)NCH experiment. The phase cycle was $\varphi_1$ = 4(y), 4(–y); $\varphi_2$ = y, –y; $\varphi_3$ = 2(x), 2(–x); $\varphi_4$ = 8(x), 8(y); $\varphi_5$ = 8(y), 8(–y); $\varphi_{rec}$ = 2(–y, y, y, –y, y, –y, –y, y), 2(y, –y, –y, y, –y, y, y, –y).

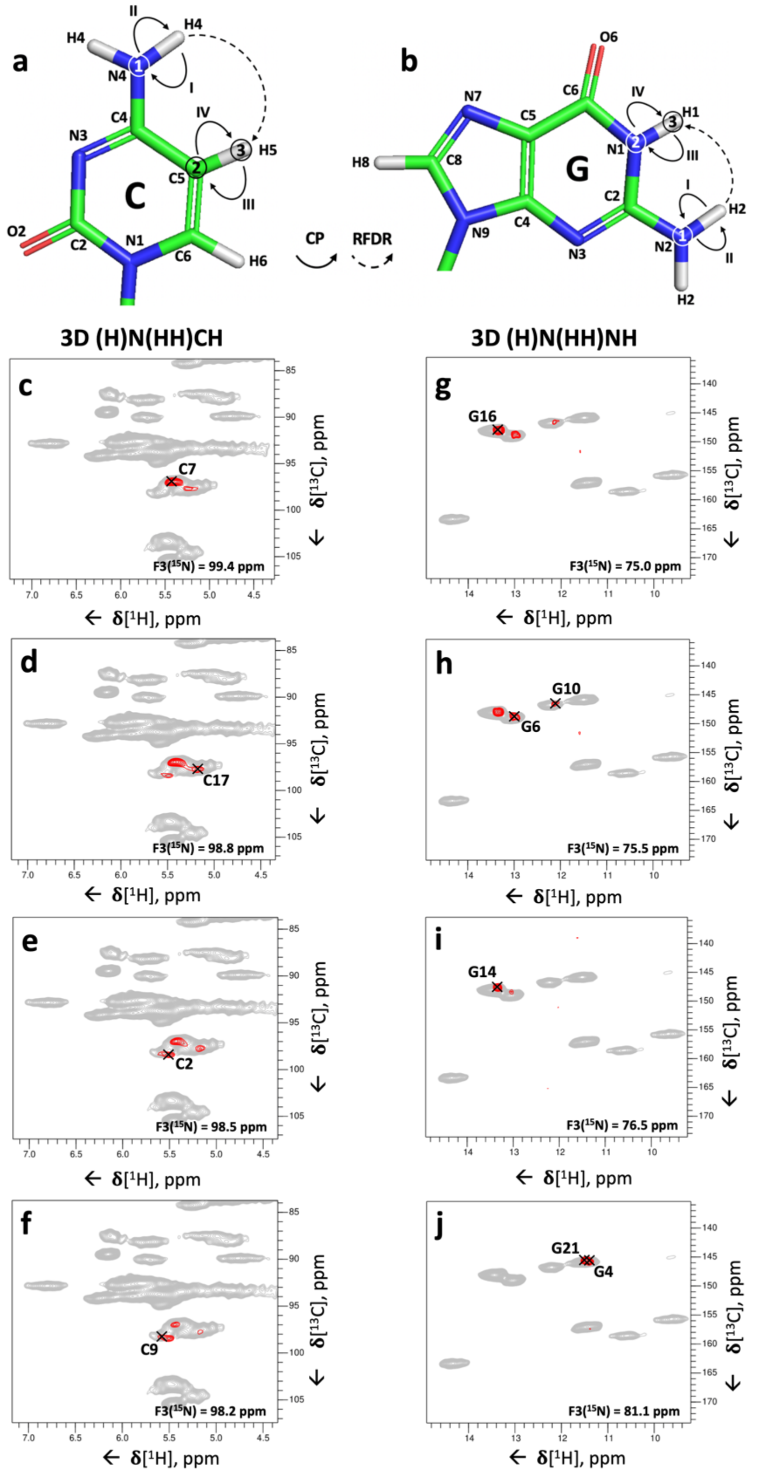


**Figure S2. 3D (H)N(HH)CH and 3D (H)N(HH)NH experiments for the assignment of nucleobase amino resonances.** (**a**) Magnetization transfer scheme of the 3D (H)N(HH)CH experiment shown for cytidine. (**b**) Magnetization transfer scheme for the N2–N1–H1 correlations of guanosines in the 3D (H)N(HH)NH experiment. Encircled numbers indicate the chemical-shift evolution times ($t_1$–$t_3$) corresponding to the three spectral dimensions; roman numerals I, II, III and IV indicate the first, second, third and fourth CP transfer periods, respectively. (**c-f**) Representative 2D $^{1}$H -$^{13}$C planes from the 3D (H)N(HH)CH spectrum showing cytidine N4–C5–H5 correlations for the nucleotides C7 (**c**), C17 (**d**), C2 (**e**) and C9 (**f**). The $^{1}$H-$^{1}$H RFDR mixing time was 0.48 ms. (**g-j**) Representative 2D $^{1}$H-$^{15}$N planes extracted from the 3D (H)N(HH)NH spectrum showing guanosine N2–N1–H1 correlations for the nucleotides G16 (**g**), G6 and G10 (**h**), G14 (**i**), G21 and G4 (**j**). The $^{1}$H-$^{1}$H RFDR mixing time was 0.48 ms. For reference, in panels (c-f) the red contours of the 3D (H)N(HH)CH spectrum are overlaid on the 2D $^{1}$H-$^{13}$C CP-HSQC spectrum (in grey) tailored for the ribose/C5-H5 spectral region; in panels (g-j) the red contours of the 3D (H)N(HH)NH spectrum are overlaid on the 2D $^{1}H$-$^{15}N$ CP-HSQC spectrum (in grey). The pulse sequences and phase cycles are given in Fig. S1a and b.

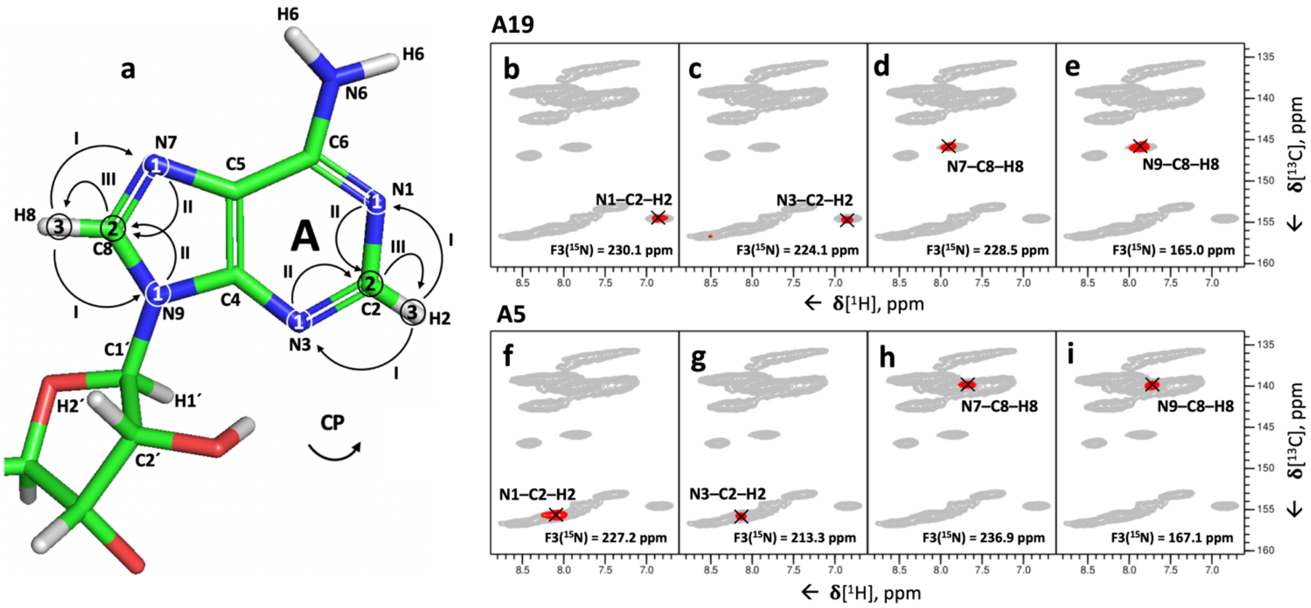


**Figure S3. Assignment of non-protonated nucleobase nitrogen chemical shifts by the 3D (H)NCH experiment.** (**a**) Magnetization transfer scheme of the 3D (H)NCH experiment shown for adenosine as an example. Encircled numbers indicate the chemical-shift evolution times ($t_1$–$t_3$) corresponding to the three spectral dimensions; roman numerals I, II and III indicate the first (5 ms), second (8 ms) and third (0.25 ms) CP transfer periods. (**b-i**) Representative 2D $^{1}H$ -$^{13}C$ planes extracted from the 3D (H)NCH spectrum showing the A19-N1–C2–H2 cross-peak (**b**), the A19-N3–C2–H2 cross-peak (**c**), the A19-N7–C8–H8 cross-peak **(d**), the A19-N9–C8–H8 cross-peak (**e**), the A5-N1–C2–H2 cross-peak (**f**), the A5-N3–C2–H2 cross-peak (**g**), the A5-N7–C8–H8 cross-peak **(h**), the A5-N9–C8–H8 cross-peak (**i**). The pulse sequence and phase cycle of the 3D (H)NCH experiment are given in Fig. S1g. For reference, in panels (b-i) the red contours of the 3D(H)N(C)CH spectrum are overlaid on the 2D $^{1}H$-$^{13}C$ CP-HSQC spectrum (in grey) tailored for nucleobase resonances.

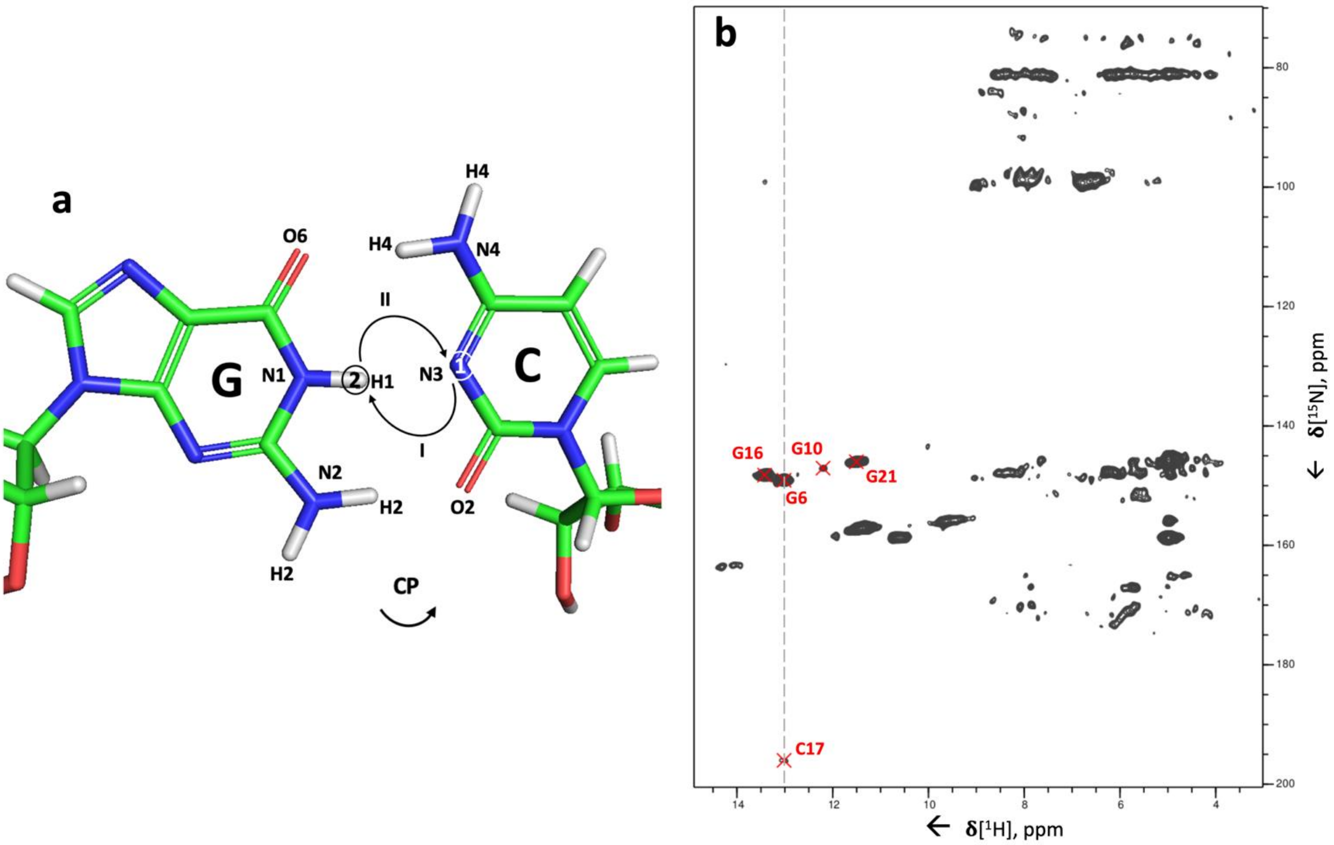


**Figure S4. Assignment of C-N3 resonances by the 2D long-range $^{1}$H-$^{15}$N CP-HSQC experiment.** (**a**) Magnetization transfer scheme of the 2D $^{1}$H-$^{15}$N CP-HSQC experiment shown for inter-strand G-H1–C-N3 correlations. Encircled numbers indicate the chemical-shift evolution times ($t_1$ & $t_2$) corresponding to the two spectral dimensions; roman numerals I and II indicate the first and second CP transfer times (both 8 ms). (**b**) 2D $^{1}$H-$^{15}$N CP-HSQC experiment with 8 ms-long $^{1}$H-$^{15}$N/$^{15}$N-$^{1}$H CP transfer times. The pulse sequence and phase cycle of the 2D $^{1}$H-$^{15}$N CP-HSQC experiment are given in Fig. S1c.

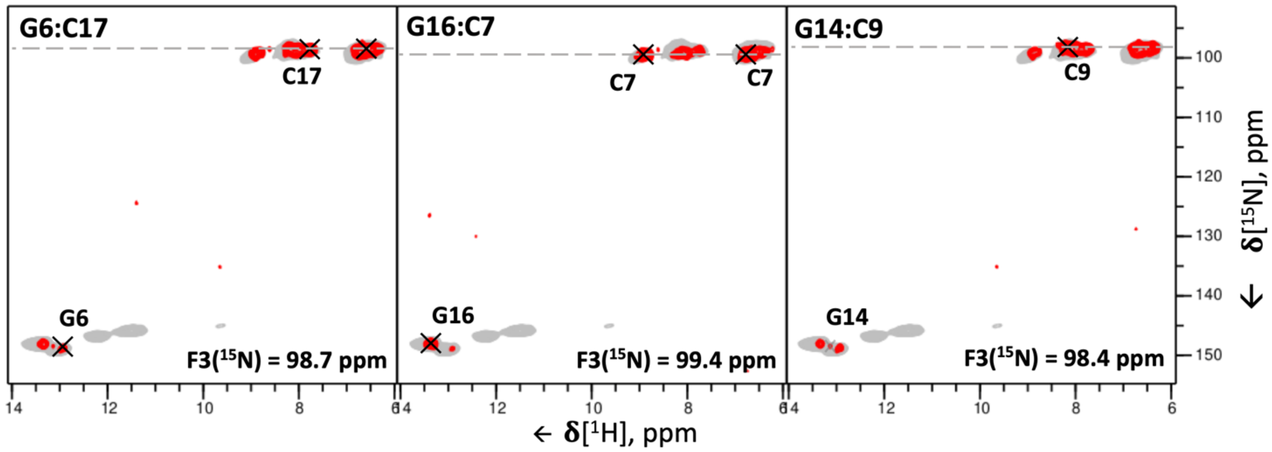


**Figure S5. Detection of G:C base pairs by the 3D (H)N(HH)NH experiment.** Representative 2D $^{1}H$-$^{15}N$ planes extracted from the 3D (H)N(HH)NH spectrum showing C-N4–G-N1–H1 cross-peaks for the base pairs G6:C17, C7:G16 and C9:G14. The pulse sequence and phase cycle of the 3D (H)N(HH)NH experiment are given in Fig. S1b. For reference, the red contours of the 3D (H)N(HH)NH spectrum are overlaid on the 2D $^{15}N$-$^{1}H$ CP-HSQC spectrum (in grey).

**Table S1.** Acquisition parameters and evolution times of all spectra.

| Spectrum ↓ | Indirect evolution times, ms | | | | | Carrier frequencies, ppm | | | Acq, ms | Spectral widths, kHz (ppm) | | | | | | RFDR mixing time, ms | Scans per point | Exp. time, h |
|---|---|---|---|---|---|---|---|---|---|---|---|---|---|---|---|---|---|---|
| **Dimensions →** | $^{1}H$ | $^{13}C$ | $^{13}C$ | $^{15}N$ | $^{15}N$ | $^{1}H$ | $^{13}C$ | $^{15}N$ | $^{1}H$ | $^{1}H$ | $^{13}C$ | $^{13}C$ | $^{15}N$ | $^{15}N$ | $^{1}H$ acq | | | |
| 2D $^{1}H$-$^{13}C$ HSQC ribose-selective | - | 6.0 | - | - | - | 7.1 | 70 | 170 | 12.8 | - | 21 (100) | - | - | - | 40 (47) | - | 32 | 5.1 |
| 2D $^{1}H$-$^{13}C$ HSQC base-selective | - | 7.5 | - | - | - | 7.1 | 140 | 170 | 12.8 | - | 13 (60) | - | - | - | 40 (47) | - | 64 | 7.6 |
| 2D $^{1}H$-$^{15}N$ HSQC | - | - | - | 9.7 | - | 7.1 | 140 | 117.5 | 12.8 | - | - | - | 10 (120) | - | 40 (47) | - | 64 | 7.9 |
| 2D $^{1}H$-$^{15}N$ HSQC, 50 % $D_2O$ | - | - | - | 9.7 | - | 7.1 | 140 | 117.5 | 12.8 | - | - | - | 10 (120) | - | 40 (47) | - | 64 | 6.2 |
| 2D $^{1}H$-$^{15}N$ HSQC, long-range | - | - | - | 8.2 | - | 7.1 | 140 | 135 | 12.8 | - | - | - | 12 (140) | - | 40 (47) | - | 138 | 16 |
| 2D (H)N(HHC)H | - | - | - | 5.8 | - | 7.1 | 160 | 160 | 12.8 | - | - | - | 13.8 (160) | - | 40 (47) | 0.40 | 128 | 12 |
| 2D (H)N(HHN)H | - | - | - | 6.2 | - | 7.1 | 140 | 117.5 | 12.8 | - | - | - | 10 (120) | - | 40 (47) | 0.48 | 192 | 15 |
| 3D (H)CCH base-selective | - | 3.7 | 3.7 | - | - | 7.1 | 135 | 170 | 10.0 | - | 17 (80) | 17 (80) | - | - | 40 (47) | 8 | 16 | 137 |
| 3D (H)CNH amino-selective | - | 4.4 | - | 3.7 | - | 7.1 | 155 | 80 | 12.8 | - | 11 (50) | - | 4.3 (50) | - | 40 (47) | - | 32 | 61 |
| 3D (H)CNH imino-selective | - | 5.6 | - | 5.4 | - | 7.1 | 160 | 154 | 12.8 | - | 4.3 (20) | - | 2.2 (26) | - | 40 (47) | - | 64 | 36 |
| 3D (H)N(C)CH base-selective | - | 2.9 | - | 3.7 | - | 7.1 | 134 | 120 | 6.4 | - | 6.8 (32) | - | 8.6 (100) | - | 40 (47) | 14 | 64 | 80 |
| 3D H(NC)CH base-selective | 2.1 | 2.9 | - | - | - | 7.1 | 130 | 120 | 6.4 | 9.4 (11) | 8.6 (40) | - | - | - | 40 (47) | 14 | 64 | 62 |
| 3D (H)NCH base-selective | - | 5.0 | - | 5.9 | - | 7.1 | 149 | 190 | 12.8 | - | 6.4 (30) | - | 9.5 (110) | - | 40 (47) | - | 16 | 56 |
| 3D (H)N(HH)CH, short RFDR | - | 4.5 | - | 5.6 | - | 7.1 | 85 | 120 | 12.8 | - | 11 (50) | - | 8.6 (100) | - | 40 (47) | 0.48 | 16 | 91 |
| 3D (H)N(HH)CH, long RFDR | - | 4.5 | - | 4.5 | - | 7.1 | 85 | 120 | 12.8 | - | 11 (50) | - | 8.6 (100) | - | 40 (47) | 0.96 | 24 | 111 |
| 3D (H)N(HH)NH | - | - | - | 6.0 | 6.0 | 7.1 | 140 | 117.5 | 12.8 | - | - | - | 8.6 (100) | 8.6 (100) | 40 (47) | 0.48 | 16 | 107 |
| 3D (HN)H(H)NH | 3.4 | - | - | 5.0 | - | 10.0 | 140 | 117.5 | 7.5 | 9.4 (11) | - | - | 8.6 (100) | - | 40 (47) | 0.48 | 32 | 85 |

**Table S2.** Summary of cross-polarization conditions.

| Spectrum ↓ | Transfer | Contact time, ms | Type | Max (average) RF strength, kHz | | RF pulse shape | |
|---|---|---|---|---|---|---|---|
| 2D $^1H$-$^{13}C$ HSQC ribose-selective | $^1H \longrightarrow {}^{13}C$ | 0.35 | DQ(n=1) | 73.6 | 30.2 | Linear ramp up ±30% | Rectangle |
| | $^{13}C \longrightarrow {}^1H$ | 0.20 | ZQ(n=1) | 129.4 | 30.2 | Linear ramp up ±30% | Rectangle |
| 2D $^1H$-$^{13}C$ HSQC base-selective | $^1H \longrightarrow {}^{13}C$ | 0.35 | DQ(n=1) | 73.6 | 30.2 | Linear ramp up ±30% | Rectangle |
| | $^{13}C \longrightarrow {}^1H$ | 0.20 | ZQ(n=1) | 129.4 | 30.2 | Linear ramp up ±30% | Rectangle |
| 2D $^1H$-$^{15}N$ HSQC | $^1H \longrightarrow {}^{15}N$ | 0.60 | DQ(n=1) | 80.0 | 25.2 | Linear ramp up ±30% | Rectangle |
| | $^{15}N \longrightarrow {}^1H$ | 0.80 | ZQ(n=1) | 126.1 | 25.2 | Linear ramp up ±30% | Rectangle |
| 2D $^1H$-$^{15}N$ HSQC, 50 % $D_2O$ | $^1H \longrightarrow {}^{15}N$ | 0.80 | DQ(n=1) | 78.6 | 25.2 | Linear ramp up ±30% | Rectangle |
| | $^{15}N \longrightarrow {}^1H$ | 0.65 | ZQ(n=1) | 127.5 | 25.2 | Linear ramp up ±30% | Rectangle |
| 2D $^1H$-$^{15}N$ HSQC, long-range | $^1H \longrightarrow {}^{15}N$ | 8.0 | DQ(n=1) | 86.2 | 25.2 | Linear ramp up ±30% | Rectangle |
| | $^{15}N \longrightarrow {}^1H$ | 8.0 | ZQ(n=1) | 127.5 | 25.2 | Linear ramp up ±30% | Rectangle |
| 2D (H)N(HHC)H | $^1H \longrightarrow {}^{15}N$ | 0.80 | DQ(n=1) | 81.0 | 25.2 | Linear ramp up ±30% | Rectangle |
| | $^{15}N \longrightarrow {}^1H$ | 0.60 | ZQ(n=1) | 119.8 | 25.2 | Linear ramp up ±30 % | Rectangle |
| | $^1H \longrightarrow {}^{13}C$ | 0.35 | DQ(n=1) | 73.0 | 30.2 | Linear ramp up ±30% | Rectangle |
| | $^{13}C \longrightarrow {}^1H$ | 0.45 | ZQ(n=1) | 128.4 | 30.2 | Linear ramp up ±30% | Rectangle |
| 2D (H)N(HHN)H | $^1H \longrightarrow {}^{15}N$ | 0.60 | DQ(n=1) | 81.2 | 25.2 | Linear ramp up ±30% | Rectangle |
| | $^{15}N \longrightarrow {}^1H$ | 0.80 | ZQ(n=1) | 135.5 | 25.2 | Linear ramp up ±30% | Rectangle |
| 3D (H)CCH base-selective | $^1H \longrightarrow {}^{13}C$ | 3.0 | DQ(n=1) | 73.0 | 30.2 | Linear ramp up ±30% | Rectangle |
| | $^{13}C \longrightarrow {}^1H$ | 0.50 | ZQ(n=1) | 128.4 | 30.2 | Linear ramp up ±30% | Rectangle |
| 3D (H)CNH amino-selective | $^1H \longrightarrow {}^{13}C$ | 3.0 | DQ(n=1) | 73.0 | 20.0 | Linear ramp up ±30% | Rectangle |
| | $^{13}C \longrightarrow {}^{15}N$ | 7.0 | DQ(n=1) | 62.4 | 39.1 | Rectangle | Tangential down ±10% |
| | $^{15}N \longrightarrow {}^1H$ | 1.0 | ZQ(n=1) | 119.8 | 25.2 | Linear ramp up ±30% | Rectangle |
| 3D (H)CNH imino-selective | $^1H \longrightarrow {}^{13}C$ | 3.0 | DQ(n=1) | 73.0 | 20.0 | Linear ramp up ±30% | Rectangle |
| | $^{13}C \longrightarrow {}^{15}N$ | 7.0 | DQ(n=1) | 62.4 | 39.1 | Rectangle | Tangential down ±10% |
| | $^{15}N \longrightarrow {}^1H$ | 1.0 | ZQ(n=1) | 119.8 | 25.2 | Linear ramp up ±30% | Rectangle |
| 3D (H)N(C)CH base-selective | $^1H \longrightarrow {}^{15}N$ | 1.0 | DQ(n=1) | 81.0 | 25.2 | Linear ramp up ±30% | Rectangle |
| | $^{15}N \longrightarrow {}^{13}C$ | 10.0 | DQ(n=1) | 39.9 | 62.4 | Tangential down ±10% | Rectangle |
| | $^{13}C \longrightarrow {}^1H$ | 0.4 | ZQ(n=1) | 128.4 | 30.2 | Linear ramp up ±30% | Rectangle |
| 3D H(NC)CH base-selective | $^1H \longrightarrow {}^{15}N$ | 0.60 | DQ(n=1) | 81.2 | 25.2 | Linear ramp up ±30% | Rectangle |
| | $^{15}N \longrightarrow {}^{13}C$ | 8.0 | DQ(n=1) | 40.8 | 62.4 | Tangential down ±10% | Rectangle |
| | $^{13}C \longrightarrow {}^1H$ | 0.20 | ZQ(n=1) | 128.7 | 30.2 | Linear ramp up ±30% | Rectangle |
| 3D (H)NCH base-selective | $^1H \longrightarrow {}^{15}N$ | 5.0 | DQ(n=1) | 73.0 | 25.2 | Linear ramp up ±30% | Rectangle |
| | $^{15}N \longrightarrow {}^{13}C$ | 8.0 | DQ(n=1) | 40.8 | 60.1 | Tangential down ±10% | Rectangle |

| | | | | | | | |
|---|---|---|---|---|---|---|---|
| | $^{13}C \longrightarrow {}^{1}H$ | 0.25 | ZQ(n=1) | 119.8 | 29.5 | Linear ramp up ±30% | Rectangle |
| 3D (H)N(HH)CH, short RFDR | $^{1}H \longrightarrow {}^{15}N$ | 0.80 | DQ(n=1) | 79.2 | 25.2 | Linear ramp up ±30% | Rectangle |
| | $^{15}N \longrightarrow {}^{1}H$ | 0.60 | ZQ(n=1) | 119.8 | 25.2 | Linear ramp up ±30% | Rectangle |
| | $^{1}H \longrightarrow {}^{13}C$ | 0.35 | DQ(n=1) | 73.0 | 30.2 | Linear ramp up ±30% | Rectangle |
| | $^{13}C \longrightarrow {}^{1}H$ | 0.45 | ZQ(n=1) | 128.4 | 30.2 | Linear ramp up ±30% | Rectangle |
| 3D (H)N(HH)CH, long RFDR | $^{1}H \longrightarrow {}^{15}N$ | 0.80 | DQ(n=1) | 79.2 | 25.2 | Linear ramp up ±30% | Rectangle |
| | $^{15}N \longrightarrow {}^{1}H$ | 0.60 | ZQ(n=1) | 119.8 | 25.2 | Linear ramp up ±30% | Rectangle |
| | $^{1}H \longrightarrow {}^{13}C$ | 0.35 | DQ(n=1) | 73.0 | 30.2 | Linear ramp up ±30% | Rectangle |
| | $^{13}C \longrightarrow {}^{1}H$ | 0.45 | ZQ(n=1) | 128.4 | 30.2 | Linear ramp up ±30% | Rectangle |
| 3D (H)N(HH)NH | $^{1}H \longrightarrow {}^{15}N$ | 0.80 | DQ(n=1) | 79.2 | 25.2 | Linear ramp up ±30% | Rectangle |
| | $^{15}N \longrightarrow {}^{1}H$ | 0.80 | ZQ(n=1) | 125.5 | 25.2 | Linear ramp up ±30% | Rectangle |
| 3D (HN)H(H)NH | $^{1}H \longrightarrow {}^{15}N$ | 0.45 | DQ(n=1) | 78.6 | 25.2 | Linear ramp up ±30% | Rectangle |
| | $^{15}N \longrightarrow {}^{1}H$ | 0.65 | ZQ(n=1) | 127.5 | 25.2 | Linear ramp up ±30% | Rectangle |

**Table S3.** Fourier processing parameters of all spectra.

| Spectrum ↓ | Digital resolution | | | | | | Window function | | | | | |
|---|---|---|---|---|---|---|---|---|---|---|---|---|
| **Dimensions →** | $^{1}$H | $^{13}$C | $^{13}$C | $^{15}$N | $^{15}$N | $^{1}$H acq | $^{1}$H | $^{13}$C | $^{13}$C | $^{15}$N | $^{15}$N | $^{1}$H acq |
| 2D $^{1}$H-$^{13}$C HSQC ribose-selective | - | 1024 | - | - | - | 8192 | - | QSINE2 | - | - | - | QSINE2 |
| 2D $^{1}$H-$^{13}$C HSQC base-selective | - | 1024 | - | - | - | 8192 | - | QSINE2 | - | - | - | QSINE2 |
| 2D $^{1}$H-$^{15}$N HSQC | - | - | - | 1024 | - | 4096 | - | - | - | QSINE2 | - | QSINE2 |
| 2D $^{1}$H-$^{15}$N HSQC, 50 % $D_2O$ | - | - | - | 1024 | - | 4096 | - | - | - | QSINE2 | - | QSINE2 |
| 2D $^{1}$H-$^{15}$N HSQC, long-range | - | - | - | 1024 | - | 4096 | - | - | - | QSINE2 | - | QSINE2 |
| 2D (H)N(HHC)H | - | - | - | 1024 | - | 8192 | - | - | - | QSINE2 | | GM -70Hz/0.2 |
| 2D (H)N(HHN)H | - | - | - | 1024 | - | 4096 | - | - | - | QSINE2 | - | GM -80Hz/0.2 |
| 3D (H)CCH base-selective | - | 256 | 256 | - | - | 4096 | - | QSINE2 | QSINE2 | - | - | GM -40Hz/0.2 |
| 3D (H)CNH amino-selective | - | 256 | - | 128 | - | 4096 | - | QSINE2 | - | GM -30Hz/0.2 | - | GM -40Hz/0.2 |
| 3D (H)CNH imino-selective | - | 256 | - | 128 | - | 4096 | - | QSINE2 | - | GM -30Hz/0.2 | - | GM -40Hz/0.2 |
| 3D (H)N(C)CH base-selective | - | 256 | 128 | 256 | - | 4096 | - | GM -30Hz/0.2 | - | QSINE2 | - | GM -40Hz/0.2 |
| 3D H(NC)CH base-tuned | 128 | 256 | - | - | - | 4096 | QSINE2 | GM -30Hz/0.2 | - | - | - | GM -40Hz/0.2 |
| 3D (H)NCH base-selective | - | 128 | - | 256 | - | 4096 | - | QSINE2 | - | GM -30Hz/0.2 | - | GM -70Hz/0.2 |
| 3D (H)N(HH)CH, short RFDR | - | 256 | - | 256 | - | 4096 | - | GM -30Hz/0.2 | - | QSINE2 | - | GM -40Hz/0.2 |
| 3D (H)N(HH)CH, long RFDR | - | 256 | - | 256 | - | 4096 | - | GM -30Hz/0.2 | - | QSINE2 | - | GM -40Hz/0.2 |
| 3D (H)N(HH)NH | - | - | - | 256 | 256 | 4096 | - | - | - | GM -30Hz/0.2 | QSINE2 | GM -40Hz/0.2 |
| 3D (HN)H(H)NH | 128 | - | - | 256 | - | 8192 | QSINE2 | - | - | GM -50Hz/0.2 | - | GM -70Hz/0.2 |

**Table S4.** Inter-strand and intra-residue distances in trans-Hoogsteen/sugar-edge G:A base pairs derived from published structures (Ahmed *et al.*, 2020[1] PDB entry 6TPH; Correll *et al.*, 1997[17] PDB entry 354D; Cate *et al.*, 1996[18] PDB entry 1GID).

| **atom distance in trans-Hoogsteen/sugar edge G:A base pairs, Å** | | | | |
|---|---|---|---|---|
| | **Inter-strand distance** | | **Intra-residue distance** | |
| **G:A base pair** | **G-H1´–A-H6** | **G-H2–A-H8** | **G-H1´–H2** | **A-H6–H8** |
| Ahmed *et al.*, 2020[1] 4G:22A | 2.6 | 3.5 | 4.2 | 5.1 |
| Ahmed *et al.*, 2020[1] 5A:21G | 2.3 | 3.0 | 4.5 | 5.1 |
| Correll *et al.*, 1997[17] 72G:104A | 2.4 | 3.3 | 4.6 | 4.9 |
| Cate *et al.*, 1996[18] 150G:153A | 2.4 | 2.3 | 4.5 | 4.9 |
| Cate *et al.*, 1996[18] 140A:163G | 2.2 | 3.4 | 4.5 | 4.9 |
| Cate *et al.*, 1996[18] 139A:164G | 2.4 | 2.9 | 4.5 | 4.9 |
| **Average** | **2.4** | **3.1** | **4.5** | **5.0** |
| **Standard deviation** | 0.1 | 0.4 | 0.1 | 0.1 |

**Table S5.** Isotropic chemical shifts of the 26mer box C/D RNA in the L7Ae–box C/D RNA complex, obtained in this work and our previous study[19] using $^{1}$H-detection. Chemical shifts measured here are highlighted in light-blue. Unassigned spins are indicated with a dash and ambiguous assignments are marked by with an asterisk.

| | Isotropic chemical shifts, ppm | | | | | | | | | | | | | | | | | | | | | | | | | | | |
|---|---|---|---|---|---|---|---|---|---|---|---|---|---|---|---|---|---|---|---|---|---|---|---|---|---|---|---|---|
| | C1' | C2 | C2' | C3' | C4 | C4' | C5 | C5' | C6 | C8 | H1 | H1' | H2 | H2' | H3 | H3' | H4 | H4' | H5 | H5´ | H6 | H8 | N1 | N2 | N3 | N4 | N6 | N7 | N9 |
| C2 | - | - | - | - | 168.2 | - | 98.2 | - | 140.7 | | | - | | - | | - | 8.0 | - | 5.5 | - | 7.8 | | 151.8 | | - | 98.5 | | | |
| U3 | 94.9 | 154.3 | 75.6 | 71.7 | 166.2 | 83.0 | 104.6 | 64.0 | 140.1 | | | 5.6 | | 4.2 | 11.4 | 4.2 | | 4.3 | 5.3 | 4.2/4.4 | 7.4 | | 145.2 | | 157.0 | | | | |
| G4 | 90.0 | 155.9 | 75.9 | 79.5 | - | 88.2 | - | 67.8 | 160.6 | 139.2 | 11.3 | 5.7 | 5.5/7.7 | 5.0 | | 4.9 | | 4.9 | | 4.0/4.4 | | 8.1 | 145.7 | 81.1 | - | | | - | 166.9 |
| A5 | 88.2 | 155.8 | 78.5 | 79.9 | - | 83.5 | 121.1 | 70.3 | 157.3 | 139.9 | | 5.7 | 8.1 | 4.5 | | 4.9 | | 4.9 | | 4.2/4.4 | 6.1/8.5* | 7.7 | 227.2 | | 213.5 | | 81.1 | 236.9 | 167.2 |
| G6 | 92.7 | 157.9 | 76.0 | 77.2 | 152.4 | 83.5 | - | 70.9 | 161.9 | 139.0 | 13.0 | 6.8 | 9.0/9.3 | 4.9 | | 4.7 | | 5.0 | | 4.5/4.8 | | 8.3 | 149.0 | 75.5 | - | | | - | 170.9 |
| C7 | 93.8 | 159.1 | 75.3 | 72.1 | 168.7 | 81.9 | 96.9 | 64.3 | 142.5 | | | 6.0 | | 4.5 | | 4.6 | 6.8/8.9 | 4.5 | 5.4 | 4.2/4.7 | 8.4 | | 152.4 | | - | 99.4 | | | |
| U8 | 93.7 | 153.0 | - | 71.8 | 169.4 | - | 103.2 | - | 142.7 | | | 5.6 | | - | 14.3 | 4.7 | | - | 5.5 | - | 8.3 | | 147.9 | | 163.3 | | | | |
| C9 | 93.7 | 158.6 | - | - | 168.3 | - | 98.0 | - | 140.2 | | | 5.7 | | - | | - | 6.5/8.2 | - | 5.6 | - | 7.4 | | 152.1 | | - | 98.2 | | | |
| G10 | 93.9 | 155.9 | 75.7 | - | 157.3 | - | 120.5 | - | 161.6 | 136.2 | 12.2 | 5.7 | 6.4/7.0 | 4.5 | | - | | - | | - | | 7.6 | 146.7 | 75.7 | - | | | - | 171.2 |
| G14 | - | 157.4 | - | - | - | - | - | - | 161.6 | 137.4 | 13.3 | - | 6.7/8.6 | - | | - | | - | | - | | 8.0 | 148.1 | - | - | | | - | - |
| A15 | 92.7 | 153.2 | 75.5 | 72.2 | - | 81.6 | - | 64.2 | 157.9 | 139.7 | | 6.0 | 7.4 | 4.5 | | 4.7 | | 4.6 | | 3.9/4.6 | 7.1 | 7.9 | 222.2 | | - | | 82.6 | - | 171.7 |
| G16 | 92.5 | 157.3 | 75.4 | 72.1 | - | 81.6 | - | 64.3 | 161.8 | 136.4 | 13.4 | 5.8 | 7.2/7.7 | 4.5 | | - | | 4.5 | | 4.1/4.6 | | 7.8 | 148.2 | 75.1 | - | | | - | 171.3 |
| C17 | 93.5 | 158.5 | 75.6 | 71.7 | 168.1 | 81.5 | 97.6 | 64.4 | 139.5 | | | 5.6 | | 4.7 | | 4.5 | 6.6/7.8 | 4.5 | 5.2 | 4.5 | 7.4 | | 150.9 | | 197.0 | 98.8 | | | |
| A18 | 94.1 | 156.7 | 76.6 | 72.6 | - | 81.6 | 118.8 | 64.6 | 158.0 | 138.9 | | 6.3 | 8.4 | 4.1 | | 4.8 | | 4.8 | | 4.4/4.6 | 6.8 | 8.1 | - | | - | | 79.7 | 231.5 | 173.2 |
| A19 | 89.8 | 154.6 | 72.2 | 78.5 | 150.8 | 85.1 | 121.8 | 67.3 | 156.5 | 146.0 | | 4.7 | 6.9 | 4.8 | | 4.6 | | 3.4 | | 3.9/4.4 | 5.9 | 7.9 | 230.2 | | 224.0 | | 76.0 | 228.5 | 164.9 |
| U20 | 89.4 | 154.3 | 75.3 | 80.5 | 168.4 | 84.2 | 105.4 | 69.5 | 147.0 | | | 6.2 | | 4.8 | 10.6 | 4.5 | | 5.3 | 5.5 | 4.0/4.1 | 8.5 | | 147.6 | | 158.5 | | | | |
| G21 | 87.4 | 156.0 | 77.5 | 80.0 | - | 87.4 | 117.9 | 68.1 | 160.5 | 139.2 | 11.5 | 6.1 | 5.3/7.6 | 5.3 | | 5.0 | | 5.2 | | 4.5 | | 8.4 | 145.8 | 81.1 | - | | | - | 168.7 |
| A22 | 92.4 | 156.6 | 75.8 | 72.3 | - | 82.9 | 118.6 | 65.8 | 157.4 | 140.1 | | 5.8 | 8.6 | 5.0 | | 4.6 | | 4.8 | | 4.2/4.6 | 6.1/8.5* | 7.9 | 223.4 | | - | | 81.2 | 231.7 | 170.3 |
| U23 | 93.0 | 151.2 | 74.6 | 72.3 | 168.0 | 82.4 | 103.7 | 62.9 | 140.5 | | | 4.9 | | 4.4 | 9.6 | 4.5 | | 4.4 | 5.4 | 4.1/4.6 | 7.6 | | 145.0 | | 155.9 | | | | |

### Author Contributions

PIA performed experiments, analysed data and wrote the paper; JK assisted in the experiments and revised the manuscript; TC designed the project, acquired funds, supervised the project and wrote the manuscript; AM designed the project, performed experiments, supervised the project and wrote the manuscript.
